# Powdery mildew exploits host plastoglobuli functions via DGAT3 and FBN2 for proliferation

**DOI:** 10.1101/2025.03.04.641535

**Authors:** Hang Xue, Catalina Kemmer, Emma H. Choi, Masakazu Iwai, Krishna K. Niyogi, Mary C. Wildermuth

## Abstract

Plastoglobuli (PG) are specialized lipid-containing compartments attached to thylakoid membranes within chloroplasts. PGs participate in various metabolic pathways in response to environmental stresses such as high light, nitrogen deficiency, and heat stress. However, their roles in biotic stress remain largely unexplored. In this study, two distinct lines of investigation converge on the importance of PGs to the powdery mildew-Arabidopsis interaction. First, powdery mildew infection results in the accumulation of PGs and PG-localized *Arabidopsis* Diacylglycerol Acyltransferase 3 (DGAT3), a triacylglycerol biosynthetic enzyme that supports powdery mildew proliferation. Second, a bioinformatic analysis led to the identification of a powdery mildew effector that is PG-localized and interacts specifically with FIBRILLIN 2 (FBN2), one of the most abundant PG core proteins. While silencing the effector limits powdery mildew proliferation, an *FBN2* knockout results in increased spore production, providing a direct link between PG function and pathogen manipulation of the host. These findings underscore the significance of PGs in host-pathogen interactions and offer new insights into the intricate mechanisms underlying the interaction of an obligate biotroph with its host.

**SIGNIFICANCE:** Plastoglobuli (PG) functions are largely unexplored in the context of host-pathogen interactions. Using the *Arabidopsis*–powdery mildew system, we uncover new roles for PGs in host-pathogen dynamics through integrated genetic, biochemical, and microscopic analyses. Mutants in PG-localized proteins impact powdery mildew proliferation. Moreover, a powdery mildew effector that interacts with the core PG protein FBN2 is identified and found to contribute to virulence. These findings suggest this obligate biotrophic pathogen exploits PGs for its own gain.

## INTRODUCTION

Chloroplasts are important hubs connecting photosynthesis and plant defense (Littlejohn et al. 2021; Liu et al. 2024; Bhattacharyya and Chakraborty 2018; Lu and Yao 2018). Beyond their well-known role in producing primary carbon-containing metabolites, chloroplasts are also involved in the synthesis of crucial defense-related molecules, including the hormones jasmonic acid (JA) and salicylic acid (SA), as well as reactive oxygen species (ROS). Infection with plant pathogens often leads to a decrease in photosynthesis as shown by chlorophyll fluorescence measurements, proteomics studies, and transcriptomics studies (Prokopová et al. 2010; Mandal et al. 2014; de Torres Zabala et al. 2015). Numerous studies have shown that changes in photosynthesis following infection can be mediated by pathogen-secreted proteins, so-called effectors, that translocate to the chloroplast, although the mechanisms by which these effectors cross membrane interfaces remain largely unknown (F. Yang et al. 2021). Many chloroplast-targeted effectors exploit the host’s protein-sorting machinery by mimicking typical chloroplast transit peptides. In *Pseudomonas syringae*, one of the best-studied phytobacteria, effectors HopI1 and HopN1 localize to chloroplasts, where they suppress SA production and reduce ROS generation, respectively (Jelenska et al. 2007; Rodríguez-Herva et al. 2012). Pathogen infections also often result in noticeable changes in chloroplast morphology, such as swollen chloroplasts, increased starch accumulation, disintegrated grana stacks, induced stromule formation, and increased plastoglobule (PG) accumulation (Bhattacharyya et al. 2015; Channarayappa et al. 1992; Otulak et al. 2015; Caplan et al. 2015). For instance, PG numbers increased more than 20-fold in plants inoculated with the necrotroph *Botrytis cinerea*, suggesting that PGs are actively involved in the plant’s response to biotic stress (Zechmann 2019).

PGs are specialized lipid-containing compartments attached to chloroplast thylakoid membranes and are involved in synthesizing and storing lipid-derived molecules, such as prenyl quinones, triacylglycerols (TAGs), fatty acid phytyl esters, and carotenoids (van Wijk and Kessler 2017). The number and clustering of PGs often change in response to stress such as high light, drought, nitrogen deprivation, and pathogens (Espinoza-Corral, Schwenkert, and Lundquist 2021; Chen et al. 1998; Gaude et al. 2007; Arzac, Fernández-Marín, and García-Plazaola 2022). Mass spectrometry analyses have identified around 30 core PG proteins, along with additional proteins recruited under specific conditions (Lundquist et al. 2012, 2013; Ytterberg, Peltier, and van Wijk 2006). Among these, the most abundant proteins are members of the plastid-specific fibrillin (FBN) family, which are thought to maintain PG structure (Singh and McNellis 2011; Torres-Romero et al. 2022). Studies have also implicated FBN involvement in disease resistance. For instance, *Arabidopsis thaliana fbn4* and *fbn1b* mutants displayed greater susceptibility to the hemibiotroph *P. syringae* (Singh et al. 2010; Cooper et al. 2003). The FBN1b ortholog in rice has also been linked to stress resistance through its interaction with SA glucosyltransferase (OsSGT1), a protein involved in SA signaling (Cooper et al. 2003). Additionally, *Arabidopsis* FBN1a, FBN1b, and FBN2 can interact with allene oxide synthase (AOS), which catalyzes an early step in the JA biosynthetic pathway and impacts JA-dependent responses (Youssef et al. 2010; Torres-Romero et al. 2022; Kim et al. 2022). Together, these initial studies suggest PGs may play a defensive role in plant-pathogen interactions. However, no targeting of a PG protein by a plant pathogen has been confirmed suggesting the impact of plant pathogens on PGs may be indirect.

Powdery mildews, obligate biotrophic fungi, represent a unique category of plant pathogens that rely entirely on their hosts for the resources necessary to complete their life cycle. Following spore germination and penetration of the plant cell host cuticle and cell wall, a feeding structure (the haustorial complex) is formed in the epidermal cell (Micali et al. 2008). Required plant nutrients are then acquired through this structure to support the production of a surface hyphal network and asexual reproduction. This results in a localized shift in plant leaf metabolism from source to sink tissue accompanied by reduction in photosynthetic gene expression, chloroplast degradation, and the formation of lipid bodies in chloroplasts of mesophyll cells underlying the epidermal cell housing the haustorial complex (Xue et al. 2024; Chandran et al. 2010). These chloroplast lipid bodies may serve as an energy source for the fungus, as lipids are transferred from the host plant to the powdery mildew during asexual reproduction and are incorporated into spore TAGs (Lee et al. 2024). *Arabidopsis* Diacylglycerol Acyltransferase 3 (DGAT3), a key enzyme responsible for synthesizing TAGs within chloroplasts, plays a crucial role in the host–powdery mildew interaction (Xue et al. 2024).

Knockout and knockdown of *DGAT3* reduces TAG levels in chloroplasts and decreases powdery mildew spore production. Although the connection between DGAT3 and PGs remains unclear, it is possible that the observed powdery mildew-induced chloroplast lipid bodies either originate from PGs or involve PG-derived components. Therefore, obligate biotrophic pathogens could target host PGs for their own gain.

Although PGs are known to play important roles in response to abiotic stresses, their functions in biotic stress, particularly in interactions with obligate biotrophs, remain largely unexplored. Therefore, in this study, we investigate the role of PGs in the powdery mildew-host interaction. We show that PGs are induced at the infection site in response to powdery mildew infection and demonstrate that DGAT3, a key contributor to powdery mildew proliferation, is localized to PGs. Additionally, we reveal that FBN1a, FBN1b, and FBN2 inhibit powdery mildew proliferation, with FBN2 being specifically targeted by a powdery mildew effector that contributes to virulence. These findings provide direct evidence for pathogen manipulation of PGs and new insights into the involvement of PGs in host-pathogen interactions.

## RESULTS

### *Arabidopsis* DGAT3 is localized to PGs

*Arabidopsis* DGAT3 is localized to the chloroplast and contributes to TAG synthesis inside the chloroplast in response to powdery mildew, concurrent with the formation of chloroplast lipid bodies (Xue et al. 2024). PGs are often induced in response to stress (Espinoza-Corral, Schwenkert, and Lundquist 2021; Chen et al. 1998; Gaude et al. 2007; Arzac, Fernández-Marín, and García-Plazaola 2022) and known to contain TAGs (van Wijk and Kessler 2017). To explore the connection between DGAT3 and PGs, we transiently co-expressed DGAT3-GFP and the PG marker FBN1b-mScarlet (Francisco M. Gámez-Arjona et al. 2014) in *Nicotiana benthamiana* leaves via *Agrobacterium* infiltration. In multiple independent experiments, FBN1b-mScarlet expression was very low; this included experiments in which the P19 silencing repressor (Norkunas et al. 2018) was transiently co-expressed with FBN1b-mScarlet. However, FBN1b-mScarlet fluorescence, when observed, colocalizes with DGAT3-GFP in PG-like punctate structures associated with chloroplasts (**Fig. 1A**).

**Figure 1.**
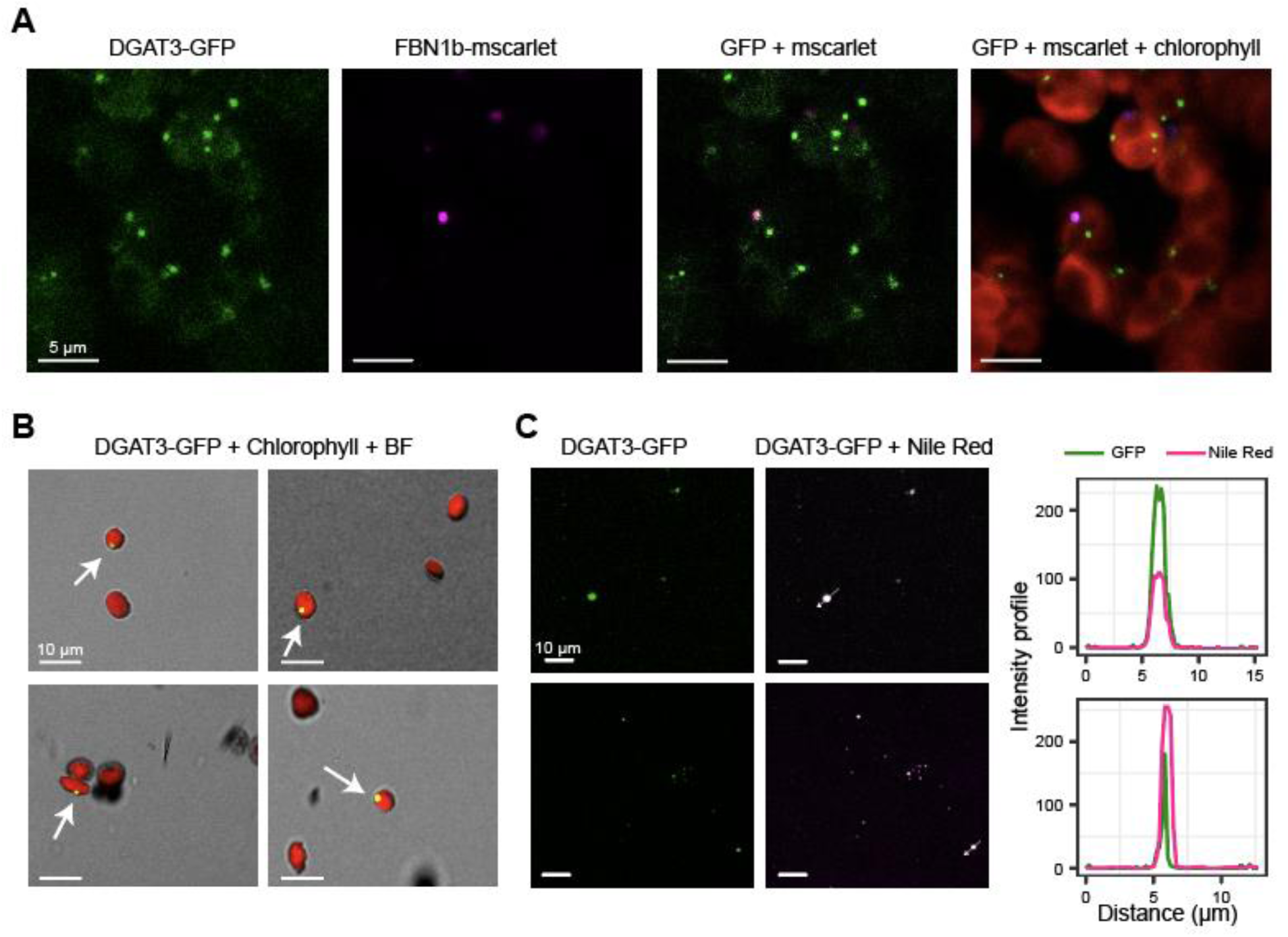
DGAT3 is localized to plastoglobule. A) Confocal microscopy images of transient co-expression of 35S:AtDGAT3-GFP and 35S:FBN1b-mScarlet in *Nicotiana benthamiana*. B) Four representative confocal microscopy images of chloroplasts isolated from 3-day darkness-incubated (3DDI) AtDGAT3-GFP transgenic *Arabidopsis* plants. C) Representative confocal microscopy images of plastoglobules isolated from 3DDI AtDGAT3-GFP transgenic *Arabidopsis* plants and stained with Nile red neutral lipid dye.

To further confirm the localization of DGAT3 to PGs, we generated a stable transgenic Arabidopsis line overexpressing DGAT3-GFP under the control of a 35S promoter. Prolonged darkness is a rapid reliable means to induce PG formation (Tominaga et al. 2018); therefore, we first treated detached leaves from these DGAT3-GFP transgenic plants with a 3-day darkness incubation (3DDI) on Murashige and Skoog agar plates at 23°C and then performed our confocal analysis on isolated chloroplasts to limit background signal. DGAT3-GFP appears as punctate dots in chloroplasts isolated from 3DDI leaves (**Fig. 1B**), with GFP fluorescence detected in approximately 15% of the ∼200 chloroplasts examined. The DGAT3 fluorescence pattern in the transgenic line is similar to that of a previously reported transgenic line expressing the PG-localized protein SOUL heme–binding protein 4 (SOUL4) (Shanmugabalaji, Grimm, and Kessler 2020). Additionally, PGs were isolated from 3DDI leaves and stained with Nile Red, a neutral lipid stain (**Fig. 1C**). DGAT3-GFP is observed in isolated PGs, confirming the PG localization of DGAT3. Furthermore, consistent with a role for DGAT3 in PG TAG synthesis, the DGAT3-GFP signals fully overlap with the Nile Red signals, as shown by pixel intensity profiling (**Fig. 1C**). Overall, our results show that DGAT3 is localized to PGs, the lipid-rich punctate structures in chloroplasts.

### Powdery mildew infection induces PG formation

Previous studies have reported an increase in the number, and sometimes the size, of PGs in response to biotic stress (Zechmann 2019; Arzac, Fernández-Marín, and García-Plazaola 2022). Several stable transgenic *Arabidopsis* lines overexpressing fluorescently labeled PG proteins are available (Shanmugabalaji, Grimm, and Kessler 2020; Rottet et al. 2016). However, none have been reported to effectively serve as markers for the dynamic changes in PGs under varying conditions. Given that DGAT3 is specifically localized to PGs and plays an important role in our interaction, we investigated whether DGAT3-GFP could serve as a reliable marker for assessing PG abundance in response to powdery mildew infection. We checked GFP fluorescence in DGAT3-GFP transgenic leaf tissues at 10 days post inoculation (10 dpi) with the powdery mildew *Golovinomyces orontii MGH1* and used untreated leaves as the negative control, and 3DDI as the positive control. While the untreated leaf tissues showed minimal GFP signals, GFP-labeled punctate structures in chloroplasts accumulate under both 10 dpi and 3DDI conditions (**Fig. 2A**). Quantification of the GFP fluorescence using a plate reader showed a ∼10-fold increase at 10 dpi and a ∼20-fold increase following 3DDI, compared to untreated controls (**Fig. 2B**). Immunoblotting of total leaf protein extracts using an anti-GFP antibody confirmed the accumulation of DGAT3-GFP at 10 dpi in response to powdery mildew infection (**Fig. 2C**).

**Figure 2.**
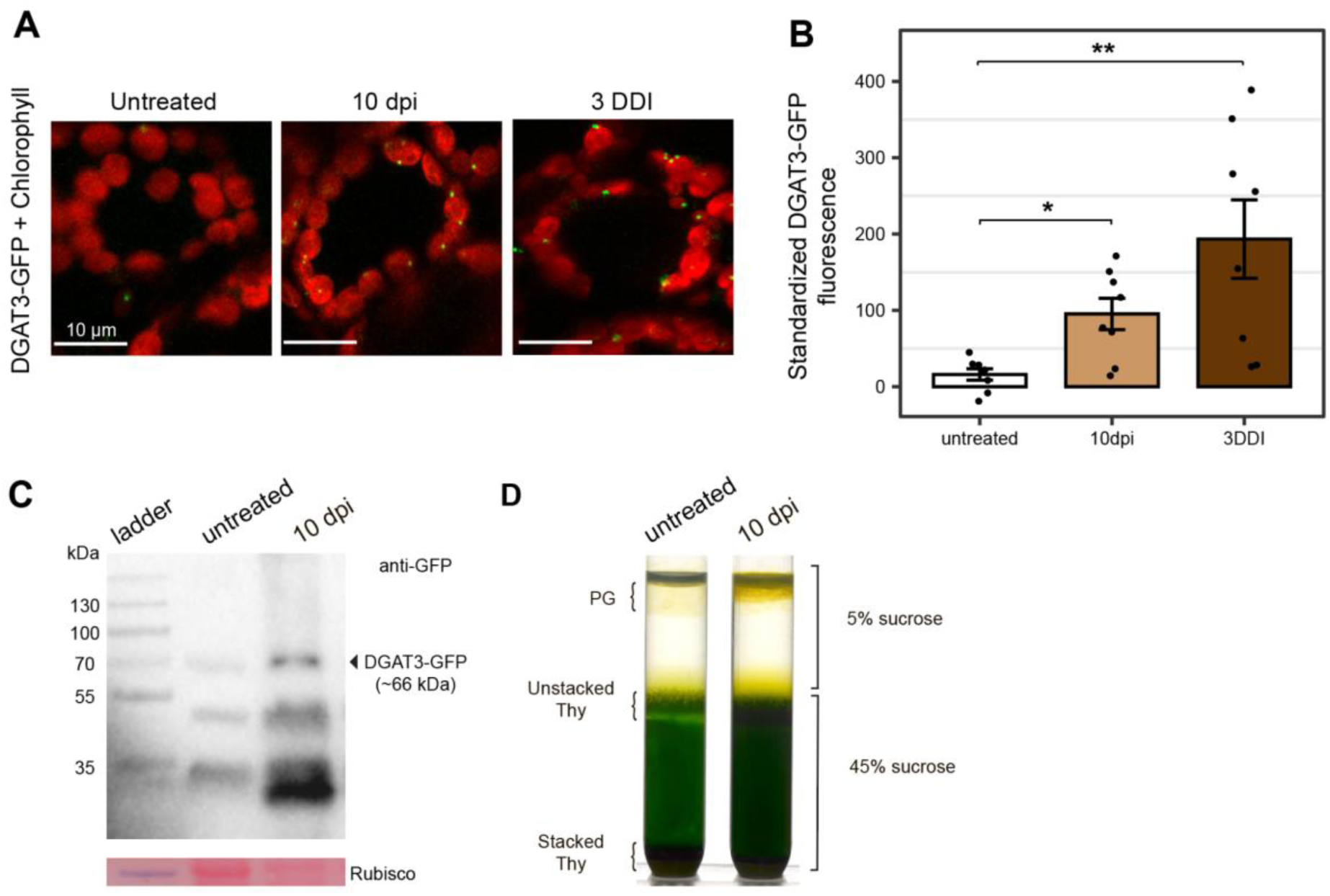
Powdery mildew infection induces DGAT3 protein accumulation and PG formation. A) Representative images of mesophyll cells from 35S:DGAT3-GFP transgenic *Arabidopsis* rosette leaves, displaying GFP signals under three conditions: untreated, 10 days post-inoculation (10 dpi) with powdery mildew, and after a 3-day dark incubation (3DDI). B) Quantification of GFP fluorescence in 35S:DGAT3-GFP transgenic *Arabidopsis* rosette leaf punches using a plate reader (mean ± SE, n = 8 leaf punches from 4 individual plants, and each data point is the mean of 8 technical replicates of fluorescence measurement). C) Proteins extracted from two leaf punches of 35S:DGAT3-GFP transgenic *Arabidopsis* plants were subjected to immunoblot analysis using the anti-GFP antibody. Ponceau S-stained membranes are shown below the blots to indicate the amount of protein loaded per lane. D) Representative images of plastoglobules separated into the top layer of a discontinuous sucrose gradient by centrifugation. An equal amount of chlorophyll (∼10 mg) for each treatment was loaded for centrifugation.

To further validate the induction of PGs by powdery mildew infection, we isolated PGs from untreated and 10 dpi wild-type Col-0 (WT) plants using discontinuous sucrose gradient centrifugation (**Fig. 2D**). The appearance of a darker yellow layer atop the 5% sucrose layer in the 10 dpi samples indicates PG accumulation during infection. Additionally, the darker green color of the unstacked thylakoid layer and the smaller pellet of stacked thylakoids observed in the 10 dpi sample indicate chloroplast dismantling (**Fig. 2D and Supplementary Figure 1)**, consistent with previous reports of chloroplast degradation during *G. orontii* infection (Xue et al. 2024). Overall, our findings demonstrate that powdery mildew infection induces PG formation and DGAT3 accumulation and that DGAT3-GFP can serve as a robust marker for tracking PG abundance.

### DGAT3, but not PG proteins PES1 and PES2, contribute to powdery mildew spore production

DGAT3 has previously been shown to contribute to powdery mildew proliferation through chloroplast TAG synthesis (Xue et al. 2024). The chloroplast PG-localized enzymes PHYTYL ESTER SYNTHASE 1 (PES1) and PES2 also have the capacity to catalyze the final acylation step in the TAG biosynthetic pathway (Lippold et al. 2012). Specifically, Lippold *et. al* showed that the *pes1 pes2* null mutant accumulates 30% less leaf TAG compared to WT plants under nitrogen-deficient conditions. To investigate whether DGAT3, PES1, and PES2 contribute synergistically to powdery mildew proliferation, we assessed spore counts on leaves of *pes1-1*, *pes2-1*, and *dgat3-2* single mutants, as well as *pes1 pes2* double mutants and *pes1 pes2 dgat3* triple mutants compared to WT plants treated in parallel. The powdery mildew spore counts in *pes1*, *pes2*, and *pes1 pes2* mutants are similar to those in WT plants (**Fig. 3**). Spore counts in the *dgat3* single mutant are significantly lower than in WT plants, as previously reported (Xue et al. 2024), as are the *pes2 dgat3* double mutant, and the *pes1 pes2 dgat3* triple mutant. Notably, the reduction in spore counts observed in the *pes2 dgat3* double mutant and *pes1 pes2 dgat3* triple mutant is not additive compared to the decrease seen in the *dgat3-2* single mutant, suggesting that DGAT3 is the primary contributor to this pathway.

**Figure 3.**
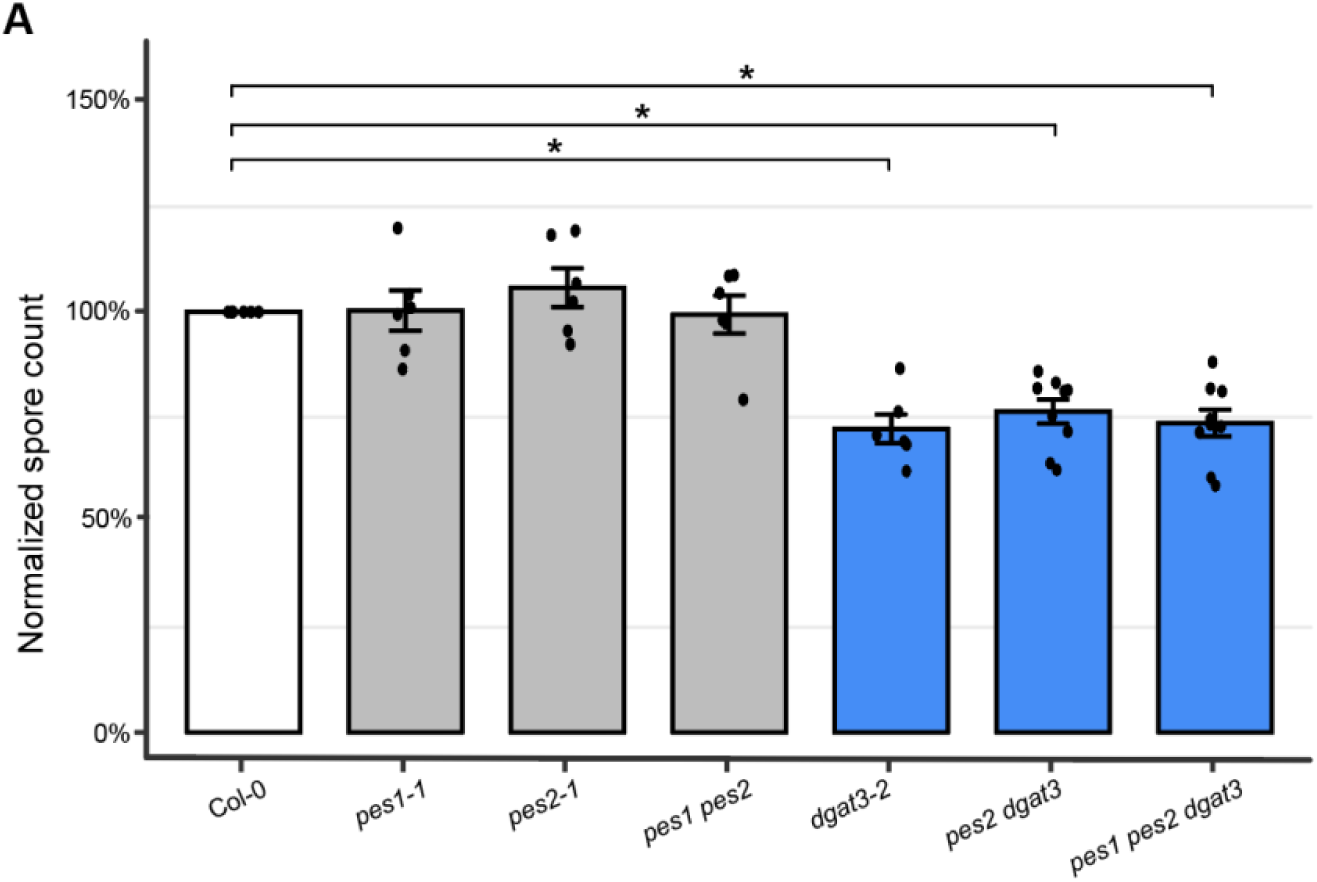
DGAT3, but not PES1 nor PES2, contribute to powdery mildew spore production. A) Spore counts/mg leaf FW at 9 dpi on mutant plants normalized to WT plants (mean ± SE, n= 6–9 independent inoculation events).Significance by unpaired, two-tailed one-sample T-test, *p < 0.05.

### Identification of a chloroplast-localized powdery mildew effector Golor4_2520209 that contributes to powdery mildew virulence

To manipulate host chloroplast functions, pathogens can direct secreted effector proteins to the host chloroplast by utilizing host-like cleavable chloroplast transit peptides (cTPs). Characterized chloroplast-targeted fungal effectors, including Pst_12806 (Xu et al. 2019) and CSEP080 (Mu et al. 2023), possess both a canonical N-terminal secretion signal and a cTP. Therefore, as shown in **Figure 4**, bioinformatics was utilized to identify candidate *G. orontii* MGH1 chloroplast-targeted effector proteins. This was followed by transient expression assays in *N. benthamiana* to identify those that are chloroplast-localized, examination of their expression over the course of infection, and functional assessment via spray-induced gene silencing (SIGS).

**Figure 4.**
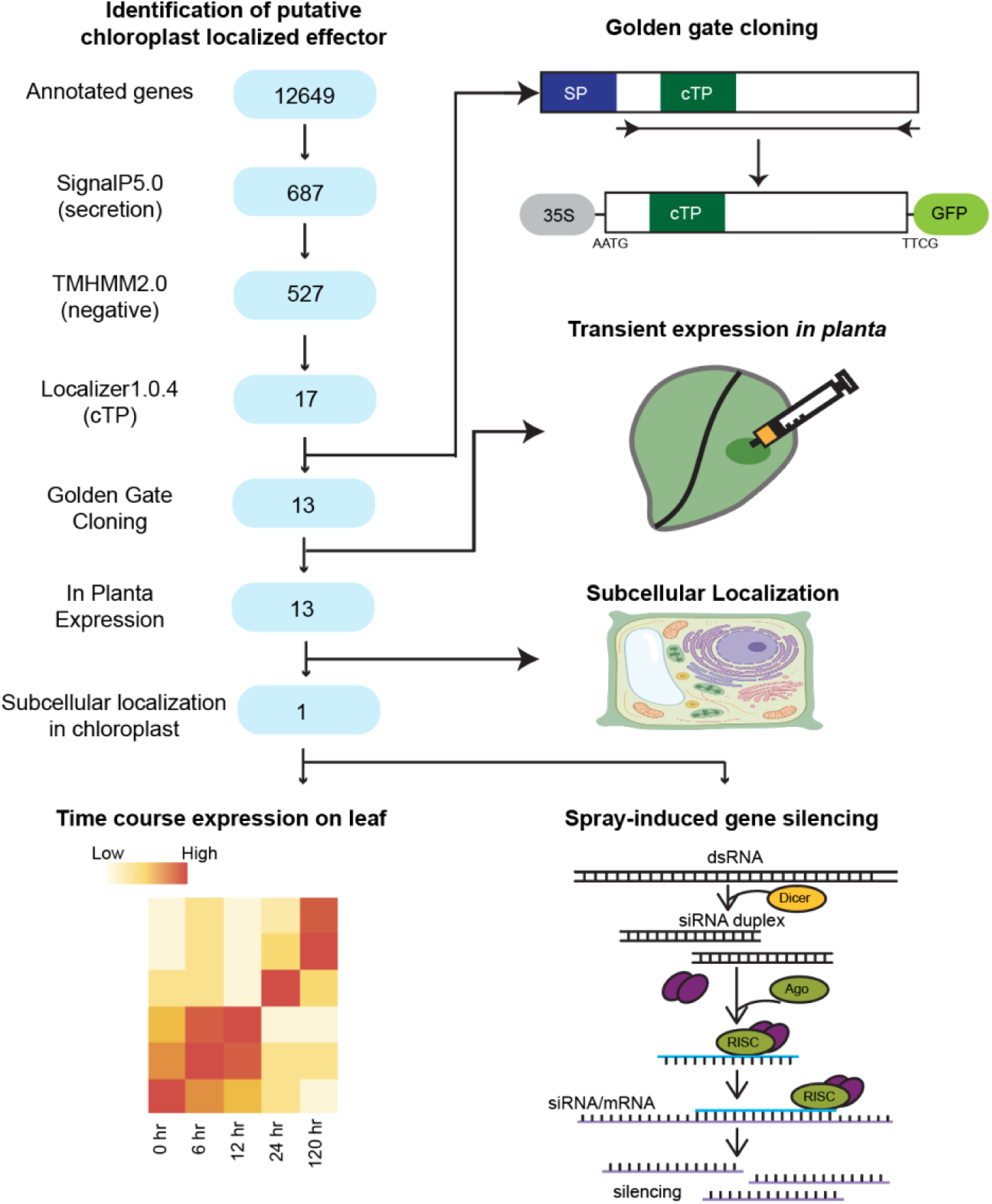
Overview of the pipeline for identifying chloroplast-localized effectors. A total of 17 candidate effectors containing a signal peptide, lacking transmembrane domains, and harboring a chloroplast transit peptide were identified from the Golor4 protein database. Of these, the coding sequences of 13 mature effectors were successfully cloned into a C-terminal GFP fusion construct under the control of a 35S promoter using Golden Gate cloning. The GFP-tagged fusion proteins were transiently expressed in *N. benthamiana* leaves through agroinfiltration. Subcellular localization in leaf pavement cells was assessed via confocal microscopy, revealing that only one effector localized specifically to the chloroplast. The expression dynamics of this effector were further analyzed using the publicly available RNA-seq dataset. The effector was transiently silenced using spray-induced gene silencing to evaluate its role in virulence.

The *G. orontii* MGH1 v4.0 genome, assembled and annotated by the Joint Genome Institute (JGI), encodes 12,649 genes. SignalP5.0 (Almagro Armenteros et al. 2019) and TMHMM2.0 (Krogh et al. 2001) were utilized to identify *G. orontii* proteins that possess a canonical secretion signal but lack any transmembrane domain(s) outside the signal peptide region. Localizer v1.0.4 was then used to predict cTPs (Sperschneider et al. 2017). In total, seventeen *G. orontii* putative chloroplast-targeted effectors (and their nearly identical homologs due to a recent whole genome duplication event) were predicted based on sequence signatures (**Supplemental Table S1**). Using Golden Gate cloning, thirteen of these putative chloroplast-targeted effectors were successfully fused at their C-terminal end to GFP for subsequent transient expression assays in *N. benthamiana*. Of these, six exhibit very low fluorescence, assessed by confocal microscopy (**Supplemental Table S1**). Of the seven remaining, four are observed in both the nucleus and cytoplasm, and two are exclusively cytoplasmic (**Supplemental Table S1**).

Only Golor4_2520209-GFP is localized to the chloroplast (**Fig. 5A and 5B**). Notably, Golor4_2520209-GFP fluorescence is observed in PG-like punctate structures associated with chloroplasts **(Fig. 5B**). Golor4_2520209 exhibits the highest expression during the initial stages of powdery mildew colonization (**Fig. 5C**), as commonly observed for *G. orontii MGH1* effectors such as EC2 (McRae et al. 2023). To assess the contribution of Golor4_2520209 to powdery mildew proliferation, we used SIGS (as in McRae et al. 2023) to transiently silence Golor4_2520209 during the course of powdery mildew infection. The dsRNA specifically targeting Golor4_2520209 was designed and sprayed onto inoculated *Arabidopsis* plants. A 30% reduction in powdery mildew spore count is observed for dsRNA-sprayed plants compared to control water-sprayed plants (**Fig. 5D)**. Together, our results show that Golor4_2520209 is localized to the chloroplast, and possibly to plastoglobuli, and contributes to powdery mildew virulence.

**Figure 5.**
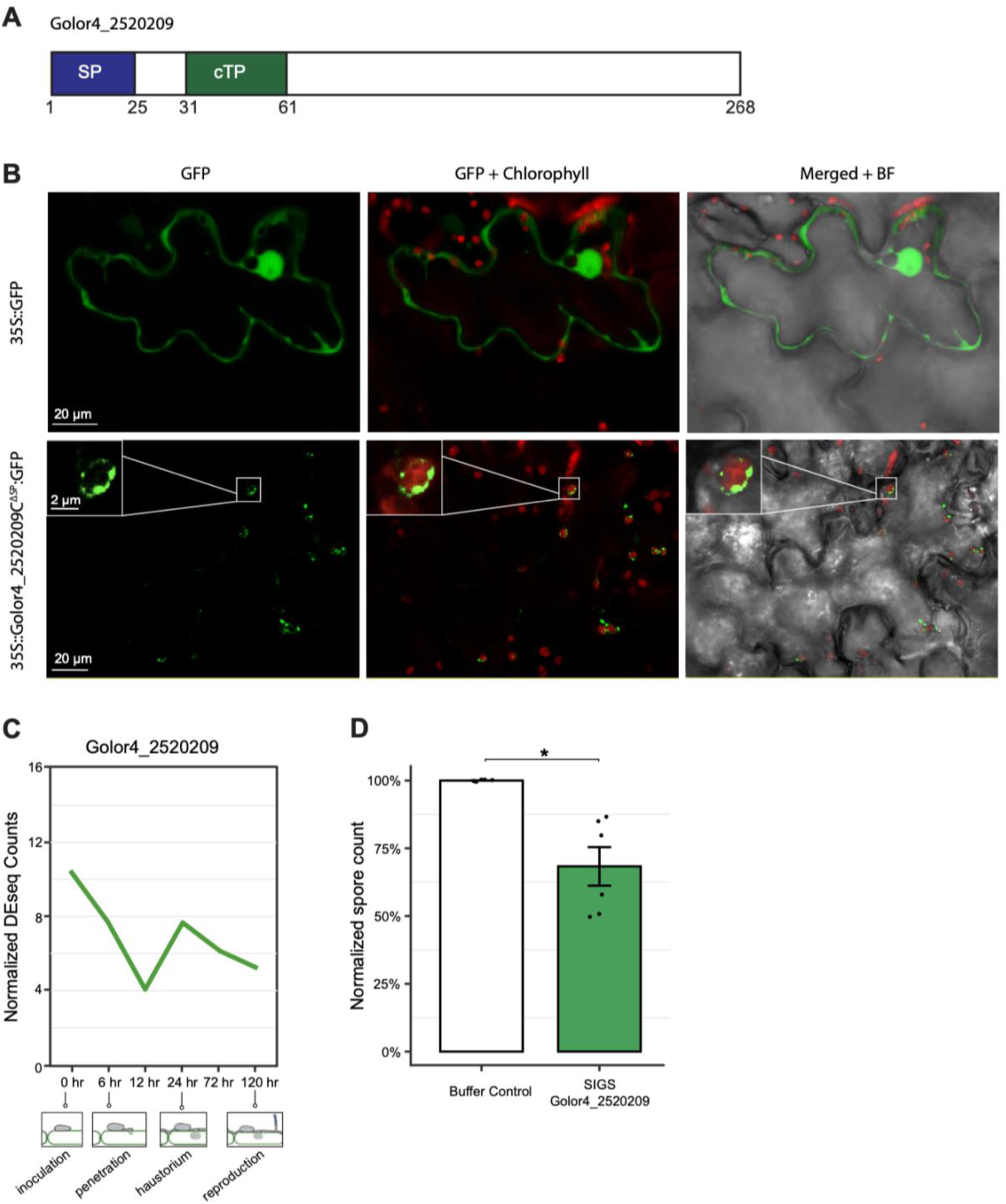
Effector Golor4_2520209 is localized to the chloroplast and contributes to powdery mildew virulence. A) Golor4_2520209 protein is predicted to have a signal peptide (1–25 aa) and a transit peptide (31–61 aa) by the SignalP5.0 and LOCALIZER 2.0 program. B) Confocal images of leaf tissues of *N. benthamiana* transiently expressing Golor4_252029ΔSP-GFP or GFP. C) Time course expression of Golor4_2520209, showing the mean of normalized DESeq2 counts from triplicate samples. D) Comparison of spore counts/mg leaf FW at 9 dpi on WT plants with Golor4_2520209 gene silenced via spray-induced gene silencing (SIGS) (± SE, n = 6 independent inoculation and silencing events).

### Golor4_2520209 interacts with the PAP_fibrillin domain of FBN2

To further explore the mechanism by which Golor4_2520209 influences chloroplast function to promote powdery mildew proliferation, we carried out a yeast two-hybrid (Y2H) screen and employed Golor4_2520209 as bait to screen the Mate & Plate™ Universal Arabidopsis library (Takara). This screen identified 12 putative interacting *Arabidopsis* proteins (**Supplementary Dataset S1**), two of which are chloroplast-localized. To validate these two proteins as putative targets of the effector Golor4_2520209, the yeast strain AH109 was co-transformed with Golor4_2520209 and the potential interactor, and grown on selective media. Of the two candidates, only the FBN2 interaction with Golor4_2520209 was confirmed (**Fig. 6A**).

**Figure 6.**
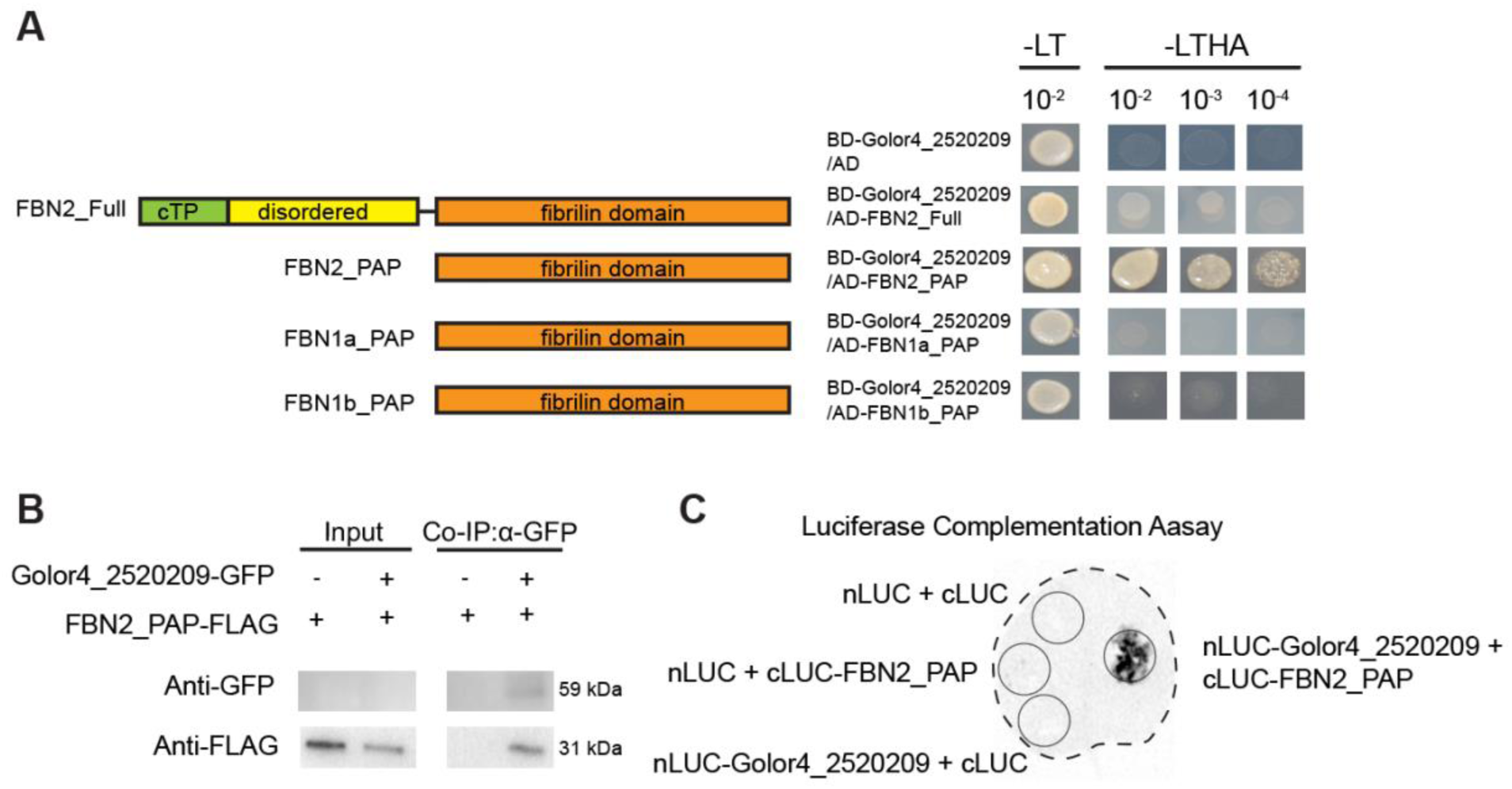
Golor4_2520209 interacts with the FBN2 fibrillin domain specifically. A) Y2H assays detect the interaction between Golor4_2520209 and the PAP_fibrilin domain of FBN2. B) Confirmation of the interaction between Golor4_2520209 and the fibrillin domain of FBN2 by co-immunoprecipitation assays. Western blots of total proteins from *N. benthamiana* leaves transiently expressing the marked constructs and proteins eluted from anti-GFP coupled magnetic beads were detected with an anti-GFP or anti-FLAG antibody. C) Confirmation of the interaction between Golor4_2520209 and the fibrillin domain of FBN2 by luciferase-complementation assay. Luciferase luminescence from *N. benthamiana* leaves transiently expressing the marked constructs was detected using a BioRad CCD imaging system.

FBN2 contains a cTP, a disordered domain, and a PAP_fibrillin domain. Our results show that Golor4_2520209 interacts specifically with the C-terminal region of FBN2 that contains the PAP_fibrillin domain (**Fig. 7A**). There is no interaction observed with the full-length FBN2 protein, possibly due to the cTP interfering with the FBN2 localization to the yeast nucleus. Of the *Arabidopsis* fibrillins, AtFBN2 is most similar to AtFBN1a/b that reside exclusively in PGs (El-Sappah et al. 2024; Francisco Manuel Gámez-Arjona et al. 2014). Therefore, we assessed whether Golor4_2520209 could interact with the PAP_fibrillin domain of FBN1a or FBN1b. Neither of these show interactions with Golor4_2520209 in yeast (**Fig. 6A**).

**Figure 7.**
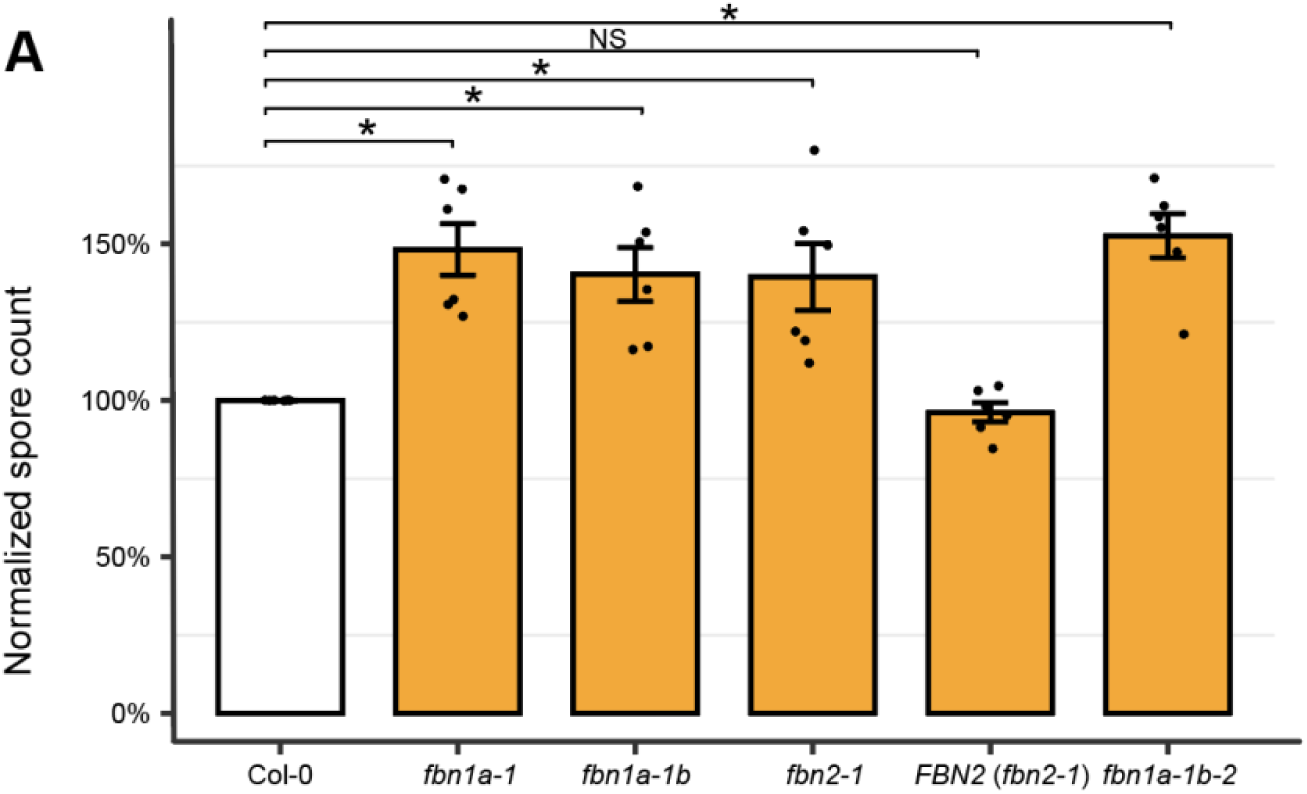
FBN1a/1b and FBN2 limit powdery mildew spore production. A) Spore counts/mg leaf FW at 9 dpi on *fbn* mutant plants and the *fbn2* complemented line normalized to WT plants (mean ± SE, n = 6 independent inoculation events). Significance by unpaired, two-tailed one-sample T-test, *p < 0.05.

We further confirmed the interaction between Golor4_2520209 and FBN2 via *in planta* co-immunoprecipitation (co-IP). Constructs expressing Golor4_2520209 tagged with a GFP and the FBN2_PAP_fibrillin domain with a FLAG-tag were transiently co-expressed in *N. benthamiana* leaves. Both Golor4_2520209 and FBN2 are detected in Western blots of proteins eluted from GFP beads (**Fig. 6B**). Additionally, the interaction was validated via *in planta* split-luciferase complementation. Golor4_2520209 and the FBN2 PAP_fibrillin domain were cloned into nLUC and cLUC vectors, respectively. When these constructs were co-expressed in *N. benthamiana* leaves, strong luminescence was observed, confirming the interaction (**Fig. 6C**). No luminescence was detected in the negative controls. In conclusion, multiple lines of evidence show the powdery mildew effector Golor4_2520209 interacts with the PG protein FBN2, indicative of direct powdery mildew manipulation of host PG function.

### FBN1a/1b and FBN2 limit powdery mildew spore production

Among the FBN family, FBN1a, FBN1b, and FBN2 are the most abundant proteins in PGs, as revealed by multiple proteomic studies (Lundquist et al. 2012, 2013; Ytterberg, Peltier, and van Wijk 2006). Furthermore, FBN1a/b and FBN2 interact with each other, are closely related, and exhibit some functional redundancy (Torres-Romero et al. 2022). However, the powdery mildew effector Golor4_2520209 specifically interacts with the PAP_fibrillin domain of FBN2 but not FBN1a/b. To distinguish their potential impact on powdery mildew infection, we assessed spore counts on leaves of the *fbn1a-1* single mutant, *fbn1a-1 fbn1b-1 (fbn1a-1b)* double mutant, *fbn2-1* single mutant, *fbn2-1* complemented line, and *fbn1a-1b fbn2* (*fbn1a-1b-2*) triple mutant. The powdery mildew spore production is significantly and similarly increased (39-48% increase) in the single, double, and triple mutants compared to WT plants, indicating not just FBN2 but FBN1 and FBN2 proteins all inhibit *G. orontii* asexual reproduction (**Fig. 7**).

Complementation of FBN2 in the *fbn2-1* mutant results in the same spore count as those measured in the WT plants, confirming the role of FBN2 in powdery mildew proliferation. Furthermore, the increased spore count in the triple mutant *fbn1a-1b-2* is not additive with respect to the increases in *fbn1a-1b* and *fbn2.* Because mutants in *FBN2* exhibit a similar impact on powdery mildew spore production to those in *FBN1* and the triple mutant, this suggests they are each required for the proper function of a shared biological process.

## DISCUSSION

### Two distinct lines of investigation converge on PGs as a key player in the powdery mildew-host interaction

PGs are specialized lipoprotein structures attached to the thylakoid membranes of chloroplasts. These dynamic compartments play key roles in the synthesis and storage of lipid-derived molecules, including prenyl quinones, TAGs, fatty acid phytyl esters, and carotenoids (van Wijk and Kessler, 2017). Typically, PGs remain sparse under normal growth conditions. However, their abundance increases significantly in response to stress (Arzac, Fernández-Marín, and García-Plazaola 2022). PGs are known to actively participate in response to abiotic stresses; however, their specific functions in host-microbe interactions remain to be explored. A role for PGs in plant defense has been inferred (e.g. through manipulation of JA biosynthesis as demonstrated in Lundquist et al. 2013 and Torres-Romero et al. 2022); however, direct targeting of PGs by pathogen effectors has not previously been demonstrated. Furthermore, previous studies did not utilize obligate biotrophs that rely on their host plant for nutrients. Herein, two independent lines of investigation converge on the important role played by PGs in the host-powdery mildew (*Arabidopsis*-*G. orontii MGH1*) interaction. Furthermore, direct targeting of the core abundant PG protein FBN2 by a powdery mildew effector that contributes to its virulence is shown.

We previously found that chloroplast-associated lipid bodies within and external to the chloroplast increase dramatically in leaf mesophyll cells underlying the epidermal cell housing the powdery mildew feeding structure concurrent with asexual reproduction (Xue et al. 2024). TAGs accumulate at the expense of thylakoid membrane lipids, with chloroplast TAGs made via DGAT3 promoting powdery mildew spore production. Transmission electron microscopy (TEM) showed some of the chloroplast lipid bodies to be associated with thylakoid membranes, a hallmark of PGs (Austin et al. 2006).

In this study, we show a marked induction of PG formation in *Arabidopsis* leaf tissues in response to powdery mildew by both differential sucrose gradient centrifugation and confocal microscopy of a labeled PG marker protein (**Fig. 2**). We also show that DGAT3 is specifically localized to the PG **(Fig. 1**) and accumulates with infection (**Fig. 2**). PG-localized proteins PES1 and PES2 also have the capacity to synthesize TAGs (Lippold et al. 2012); therefore, we explored whether they also promote powdery mildew spore production. As shown in **Figure 3**, PES1 and PES2 do not alter powdery mildew spore production nor the impact of DGAT3 on spore production. The dominant activity of PES1 and PES2 is as phytyl ester synthases that convert toxic phytol from chlorophyll breakdown to phytol esters (Lippold et al., 2012). In response to nitrogen deprivation, leaf chlorophyll breakdown is dramatic (∼18-fold), and fatty acid phytyl esters increase (Lippold et al., 2012). However, minimal reduction of chlorophyll is observed in our system, similar to the 1.5-fold reduction reported for powdery mildew infection of cucumber (Zhang, Zhou, and Wang 2022).

Similar to other plant pathogens, powdery mildews deploy effector proteins that target plant proteins, including those in the chloroplast. For instance, an effector from the grape powdery mildew, *Erysiphe necator*, interacts with the grapevine chloroplast protein VviB6f (cytochrome *b*6*f* complex iron-sulfur subunit), thereby affecting photophosphorylation and photosynthesis (Mu et al. 2023). Another study revealed that a rubber tree powdery mildew effector localizes to chloroplasts and manipulates abscisic acid biosynthesis to facilitate successful infection (X. Li et al. 2020). Therefore, we independently sought to identify powdery mildew effectors that manipulate chloroplast processes. Our bioinformatic analyses identified seventeen such potential effectors as they contain a secretion signal peptide and putative cTP (**Fig. 4, Supplemental Table S1**). *Agrobacterium*-mediated transient transformation of *N. benthamiana* with effector candidates fused to a GFP resulted in the reliable subcellular localization of seven candidates, only one of which was chloroplast-localized. The low accuracy observed in predicting effector chloroplast localization in our experiments may stem from the reliance of Localizer v1.0.4 on plant protein datasets for training, which likely do not capture the unique characteristics of effector proteins (Sperschneider et al. 2017).

Golor4_2520209 was found to be chloroplast-localized and to be associated with punctate structures indicative of PGs (**Fig. 5**). Y2H screening of an *Arabidopsis* cDNA library with Golor4_2520209 as the bait identified the core PG protein FBN2 as an interactor (**Fig. 6**). Additional Y2H studies showed Golor4_250209 interacts with the PAP_fibrillin domain of FBN2 and is specific to FBN2, not FBN1a nor FBN1b. The Golor4_250209 interaction with the PAP_fibrillin domain of FBN2 was then confirmed by two *in planta* assays that make use of *Agrobacterium*-mediated transient expression in *N. benthamiana* (**Fig. 6**): co-IP and split-luciferase complementation. SIGS of *Golor4_2520209* resulted in reduced powdery mildew spore production compared to the control treatment (**Fig. 6),** highlighting the importance of this single effector to virulence. Finally, we examined the contribution of FBN2 and the closely related FBN1 proteins to powdery mildew asexual reproduction. FBN1a/b and FBN2 are highly abundant core PG proteins that interact with each other and are postulated to form an interconnected network around PGs (Torres-Romero et al. 2022). In addition, they can specifically interact with PG-associated proteins, including the early JA biosynthetic enzyme allene oxide synthase (AOS). Disruption of *FBN1a*, *FBN1b*, and *FBN2*, singly or in combination, results in a similar increase in spore counts, suggesting they act in the same functional process, with each required for proper function (**Fig. 6**). Interestingly, a high throughput Y2H screen identified *Arabidopsis* FBN1a as a potential interactor with a powdery mildew effector candidate OE78, along with 15 other *Arabidopsis* proteins (Weßling et al. 2014). There is no OE78 ortholog in *G. orontii MGH1*, but their finding suggests that other powdery mildews may also have effectors targeting FBN1/2 proteins to manipulate PG structure and function, highlighting the importance of PGs to powdery mildew infection.

### Potential roles for PGs in the powdery mildew-host interaction

Figure 8 presents a simplified model highlighting possible mechanisms by which the powdery mildew could exploit host lipid metabolism and PG-associated proteins to facilitate its proliferation. Concurrent with its asexual reproduction, the powdery mildew induces host TAG synthesis and acquires host lipids, which are incorporated into the lipid bodies of newly formed spores (Y. Jiang et al. 2017; Lee et al. 2024). Knockout of *DGAT3* led to reduced chloroplast TAG levels and decreased spore production (Y. Jiang et al. 2017; Lee et al. 2024). Therefore, it is possible that TAGs synthesized by DGAT3 in PGs are directly acquired by the powdery mildew to support asexual reproduction.

**Figure 8.**
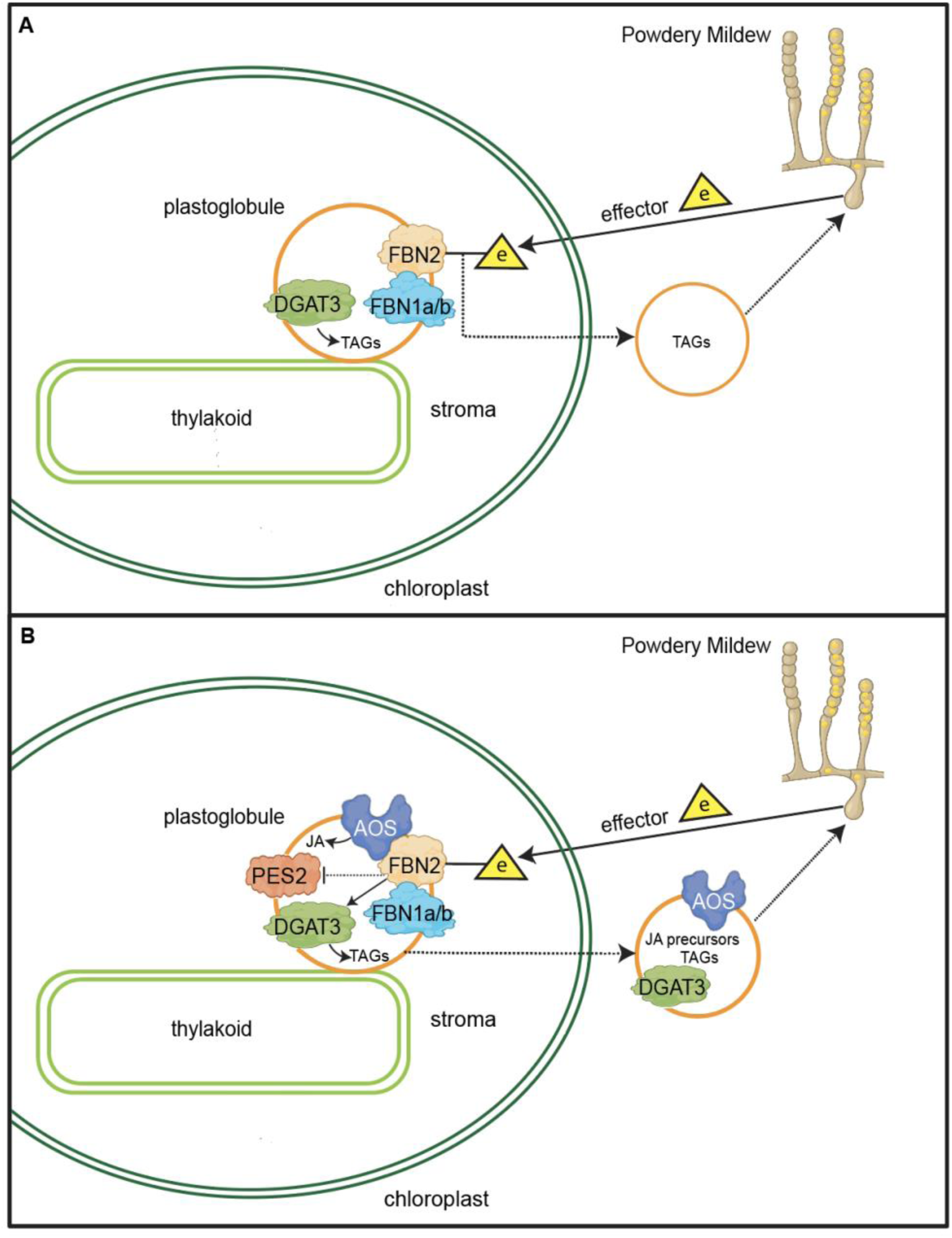
Simplified models for host PG manipulation by powdery mildew. A) Infected *Arabidopsis* leaves have an increased abundance of PGs and TAGs. DGAT3 is located in PGs and is dominantly responsible for TAG synthesis in PG. DGAT3 function benefits the fungus, likely by supplying energy dense lipids for asexual reproduction. In contrast, FBN1a/1b and FBN2 inhibit powdery mildew spore reproduction as assessed through genetic analyses. FBN1a/1b and FBN2 proteins interact, influencing PG structure and composition. The interaction between the powdery mildew effector Golor4_2520209 and FBN2 may enhance powdery mildew spore production by altering PG ultrastructure, promoting blebbing from the chloroplast, and potentially facilitating fungal access to lipids. B) Additionally, Golor4_2520209 could increase the association of AOS with PGs via FBN2. This interaction may promote the blebbing of PGs enriched in JA precursors and enzymes from the chloroplast, making them more accessible for fungal exploitation. Furthermore, Golor4_2520209 has the potential to modify the interaction between FBN2 and PES2, which could favor the stabilization and activity of DGAT3. This modification may lead to enhanced TAG production by DGAT3, further supporting fungal proliferation during infection. Abbreviations: AOS, allene oxide synthase; DGAT, diacylglycerol acyltransferase; FBN, fibrillin; JA, jasmonic acid; PES, phytyl ester synthase; PG, plastoglobule; TAGs, triacylglycerols. Dashed lines = proposed.

Stable *Arabidopsis* RNAi lines silencing the *FBN1/2* subgroup (*FBN1a*, *FBN1b*, and *FBN2*) showed no difference in chloroplast ultrastructure or PGs under unstressed conditions but altered chloroplast ultrastructure with fewer PGs in response to cold-light stress (Youssef et al. 2010). In response to powdery mildew, we found a quantitative increase in chloroplast-associated cytoplasmic globules in mesophyll cells underlying the feeding structure via TEM (Xue et al. 2024). Therefore, the interaction between the effector Golor4_2520209 and FBN2 could potentially facilitate PG blebbing from the chloroplast at the infection site, promoting *G. orontii* access to the TAGs stored within (Fig. 8A). This is likely to result from the perturbation of FBN2 propagated through the FBN2-FBN1a/b PG network and not from direct interference with the interaction of FBN2 with FBN1a/b, which appear to interact via their N-termini rather than their PAP_fibrillin domains (J. Li et al. 2020; Francisco Manuel Gámez-Arjona et al. 2014).

In addition to FBN1a and FBN1b, FBN2 has the potential to interact with other PG proteins, including AOS, and PES2 (Torres-Romero et al. 2022). Therefore, Golor4_2520209 might alter the interaction of FBN2 with one or more of these proteins, affecting PG composition, metabolism, and functionality. The best-studied interaction is that of FBN2 (and FBN1a/b) with AOS (Torres-Romero et al. 2022). If Golor4_2520209 inhibits the interaction of FBN2 with AOS, JA synthesis and/or JA-induced defense compound accumulation could be reduced, as seen in the cold- and light-stressed *FBN1A-1B-2* silenced lines (Youssef et al. 2010) or single, double, and triple *FBN1/2* mutant lines (Torres-Romero et al. 2022). Notably, the combination of cold and light stress resulted in similarly reduced levels of (JA-associated) anthocyanin accumulation for the single, double, and triple *FBN1/2* mutants (Torres-Romero et al. 2022). We also observe a similar impact (increase) on powdery mildew spore production for the single, double, and triple *FBN1/2* lines (Fig. 7). By contrast, the impact of these mutations on photosynthesis assessed as maximum PSII quantum yield (Fv/Fm) after one 1 week of cold/light stress was additive. Our data therefore suggests the impact of effector Golor4_2520209 on FBN2 would propagate through the FBN1a/b-FBN2 interconnected network. However, unlike necrotrophic pathogens, JA-associated host defense does not play a major role in limiting powdery mildew proliferation (Reuber et al. 1998), and therefore, its limitation alone is unlikely to account for the observed increases in spore production.

Instead, JA precursors and enzymes in the PG could potentially be acquired by the powdery mildew and used for its own purposes. *G. orontii* MGH1 does not have the capacity to synthesize early metabolites in JA synthesis produced in the plastid but has the predicted enzymes to catalyze subsequent reactions to JA (mycocosm.jgi.doe.gov/Golor4). In this scenario, Golor4_2520209 could increase the association of early JA biosynthetic enzymes like AOS with PGs via FBN2, and in so doing also perturb the FBN1a/b-FBN2 network facilitating blebbing of PGs enriched in JA precursors and enzymes out of the chloroplast for fungal acquisition. In addition, Golor4_2520209 has the potential to alter the interaction of FBN2 with PES2, which could result in the preferred stabilization/utilization of DGAT3 and enhanced TAG production via DGAT3. *PES1* and *PES2* single and double knockouts had no impact on powdery mildew spore production (Fig. 7), which could be explained if intact Golor4_2520209 function limits their functionality.

As FBN2 has the capacity to interact with multiple PG-localized enzymes, including FBN1a/b, AOS, and PES2, Figure 8B presents a multifunctional role for Golor4_2520209. Golor4_2520209 is secreted via the powdery mildew haustorium, taken up by the host cell (Wang et al. 2023), then translocated to chloroplast PGs that are induced in response to powdery mildew infection. Golor4_2520209 interaction with FBN2 promotes its interaction and recruitment/stabilization of AOS in PGs while limiting that of PES2, thereby promoting DGAT3 recruitment, stability, and activity. PGs would then bleb out of the chloroplast at an increased rate due to perturbation of the FBN1a/b-FBN2 network by the interaction with Golor4_2520209. Fungal acquisition of these PG-derived proteins and metabolites would thereby promote powdery mildew spore production.

### Detection and regulation of PG proteins

The proteome of *Arabidopsis* PG proteins in unstressed and light-stressed leaves has been identified by using mass spectrometry (Ytterberg, Peltier, and van Wijk 2006; Lundquist et al. 2012; Espinoza-Corral, Schwenkert, and Lundquist 2021). Interestingly, some proteins are only detectable or recruited to PGs under specific stress conditions. For instance, pheophytin pheophorbide hydrolase (PPH; AT5G13800) was identified in PG preparations from light-stressed double null mutants in *Activity of BC1 Complex Kinase 1* and *3* but not WT plants (Lundquist et al. 2013); PPH was not detectable in total leaf extracts from WT or *abc1k1-k3* stressed plants indicative of low protein abundance. DGAT3 has not been detected in previous proteomic studies of PGs or thylakoid membranes (Espinoza-Corral, Schwenkert, and Lundquist 2021; Bhuiyan et al. 2016; Froehlich et al. 2003). This absence may be due to DGAT3’s low abundance under non-stress conditions or its specific recruitment to PGs or stabilization in response to particular stressors. Our findings suggest that DGAT3-GFP levels are indeed very low in unstressed leaves but increase significantly during powdery mildew infection or darkness-induced senescence (Fig. 2A**-C**). This indicates DGAT3 may be specifically recruited to or stabilized in PGs in response to these conditions, which likely explains its absence in previous proteomic studies focused on unstressed or differently stressed tissues.

The regulation of PG proteins has been demonstrated at various levels. At the transcriptional level, for example, the transcription of *FBNs* and tocopherol cyclase (*VTE1*) is significantly increased in response to cold, drought, or high light stress (Laizet et al. 2004; Kanwischer et al. 2005). Additionally, multiple lines of evidence point to post-transcriptional and post-translational regulatory mechanisms for PG proteins. For instance, although the transcript levels of *FBN1a* were found to be comparable between the *abi2* mutant (which carries a mutation in the ABA response regulator ABI2) and WT plants, the *abi2* mutant exhibited more than 4-fold higher levels of FBN1a protein (Y. Yang et al. 2006). Stress-induced mobilization of FBN2 from the stroma to thylakoids supports the idea that post-import mechanisms can also regulate specific FBNs (Rey et al. 2000). FBN2 might then preferentially facilitate the recruitment of interacting proteins to PGs. Some FBNs, including FBN2, contain phosphorylation sites (Lohscheider, Friso, and van Wijk 2016). However, Torres-Romero et al. deemed it unlikely that FBN2 phosphorylation dictates subcellular localization (Torres-Romero et al. 2022), and Golor4_2520209 interacts with the PAP_fibrillin domain, which does not harbor any phosphorylation sites.

Our study provides additional evidence for post-transcriptional and/or post-translational regulation of the PG protein DGAT3. Microarray and RNA-seq data show that the DGAT3 transcript is not significantly induced in powdery mildew-infected *Arabidopsis* tissues at 5 dpi at the infection site or in whole leaves (Chandran et al. 2010, 2009). However, the protein level of DGAT3, expressed under the control of a constitutive promoter, is much higher in powdery mildew-infected tissues compared to uninfected controls (Fig. 2A**-C**). We only observe DGAT3-GFP localized in PGs formed in response to powdery mildew or dark induction (Fig. 2A). This could be due to the recruitment of DGAT3 to PGs in response to stress and/or the stabilization of the enzyme under these conditions. DGAT3 is unusual in that it is a [2Fe-2S] metalloenzyme that is unstable in its reduced form (Aymé et al. 2018). Thus, localized PG redox status could have a profound influence on DGAT3 stability, and a number of the FBN2 interactors identified by co-IP have redox functions (Torres-Romero et al. 2022). Therefore, Golor4_2520209 could also promote the recruitment and stability of DGAT3 through its influence on FBN2 interactors.

Together, these findings illustrate the complex, multi-layered regulation of PG proteins that allow plants to dynamically respond to environmental stress and optimize their metabolic processes accordingly.

This complex regulation of PG-associated proteins partly explains the scarcity of PG marker lines reported in the literature. Our generation of the DGAT3-GFP transgenic line offers a high-throughput and less invasive alternative to TEM or chloroplast fractionation for visualizing and quantifying PGs in live plant tissues. We used this line to monitor changes in PG formation in response to powdery mildew infection (Fig. 2A, B). By tracking GFP fluorescence, this line can be used in future studies to investigate whether the pathogen directly acquires PGs, as well as the real-time movement and clustering of PGs under other stress conditions.

Understanding how the powdery mildew manipulates and hijacks PG function for its own gain allows for unique insights into PG ultrastructure, composition, and function. Furthermore, pathogen effector targets identify key host control points (Weßling et al. 2014). Since PGs are critical for lipid metabolism, targeting them through synthetic biology may offer new avenues for engineering lipid pathways and pigment production. In addition, modulating PG protein expression can regulate cell growth metabolism in microalgae (E.-Y. Jiang et al. 2023).

Understanding the regulation of PG proteins and the strategies employed by powdery mildews to manipulate host PGs could therefore contribute to the development of new bioengineering strategies using PGs as the target sites.

## EXPERIMENTAL PROCEDURES

### Plant lines

The *Arabidopsis thaliana* mutant lines *fbn1a-1*(SALK_024528C, (Francisco M. Gámez-Arjona et al. 2014)), *fbn2-1* (SALK_124590, (Torres-Romero et al. 2022)), *pes1-1* (SALK_034549, (Lippold et al. 2012)) and *pes2-1* (SALK_071769, (Lippold et al. 2012)), *dgat3-2* (SALK_112303, (Xue et al. 2024)) were obtained from the Arabidopsis Biological Research Center. Seeds of *pes1 pes2* double mutant were obtained from Dr. Peter Dörmann (Lippold et al. 2012). Seeds of *fbn1a fbn1b* double mutant, *fbn1a fbn1b fbn2* triple mutant and the *fbn2-1* rescue line were obtained from Dr. Ángel Mérida (Torres-Romero et al. 2022). The *pes2 dgat3* double mutant was generated in this study by crossing the *pes2-1* mutant to the *dgat3-2* mutant. The *pes1 pes2 dgat3* triple mutant was generated in this study by crossing the *pes1 pes2* double mutant to the *dgat3-2* mutant. F3 homozygous progenies were used for phenotypic analyses. All mutants are in Columbia-0 (Col-0) ecotype. The previously described pICSL22010-DGAT3 construct (Xue et al. 2024), harboring 35S:DGAT3-GFP and an expression cassette of OLE1– TagRFP driven by the native OLE1 promoter, were introduced into Agrobacterium and transformed into wild type Col-0 by floral dip (Clough and Bent 1998). The 35S:DGAT3-GFP transgenic plants were selected based on RFP expression in seeds and GFP expression in leaves.

The inserted GFP sequences were verified by PCR. Homozygous stable transgenic lines were used for phenotyping. All mutants are in the ecotype Col-0 background. Genomic DNA for genotyping was extracted from leaf tissues using a CTAB based method, and homozygosity was verified through PCR with primers listed in **Supplemental Table S1**.

### Plant growth, powdery mildew spore counting and SIGS assay

*Arabidopsis thaliana* ecotype Col-0 plants were grown in Soil mix in growth chambers at 22°C, 70% relative humidity, and a 12-hour photoperiod with PAR of ∼120 μmol photons m^-2^ s^-1^. Seeds were stratified at 4C for 2-3 days before planting. For mutant phenotyping, Col-0 and mutant plants were alternately grown in 16.6 x 12.4 x 5.8 cm insert boxes, with 12 plants per box. For spray-induced gene silencing (SIGS) assays, plants were cultivated in 10.16 x 10.16 cm pots, with 4–5 plants per pot. Boxes of plants were inoculated with *Golovinomyces orontii* isolate MGH1 conidia from 4–5 fully infected leaves at 10-14 days post-inoculation (dpi) using a settling tower (Reuber et al., 1998). SIGS protocol was previously described by McRae *et al*. (2023) (McRae et al. 2023). Briefly, pssRNAit (https://plantgrn.noble.org/pssRNAit/) was used to design the dsRNA. The dsRNA was specifically designed to target and silence the gene of interest, as well as its highly similar homolog. The DNA template was amplified via PCR primers listed in **Supplemental Table S1.** The dsRNA was prepared with the HiScribe T7 High Yield RNA Synthesis Kit (New England Biolabs, Ipswich, MA, USA) and purified using the Monarch RNA Cleanup Kit (New England Biolabs, Ipswich, MA, USA). The control and dsRNA-treated plants were inoculated in the same tower to ensure equal inoculation for each replicate. Powdery mildew spore counting protocol was previously described (Xue et al. 2024). Spore counts per mg leaf fresh weight of mutant plants or dsRNA-treated plants were normalized to WT or mock counts, respectively. To determine significance, a two-tailed One-Sample T-test was performed on counts from at least 6 boxes (*p* < 0.05).

### Chloroplast and PG isolation

Leaf tissue collected from about 4–5 week old *Arabidopsis* plants under untreated, or 3-day darkness incubated (3DDI), or 12 dpi conditions were immediately homogenized by blending for 3x5 s in isolation buffer (30 mM HEPES-KOH, pH 8.0, 0.33 M sorbitol, 5 mM MgCl2, 0.1 % [w/v] BSA). The resulting homogenate was briefly filtered through one layer of Miracloth (Chicopee Mills Inc., Milltown, NJ, USA) Chloroplasts were pelleted with 5-min centrifugation at 1500 x *g* and 4°C, and washed twice with washing buffer (30 mM HEPES-KOH, pH 8.0, 0.33 M sorbitol). For confocal imaging, washed chloroplasts (pulled from 4–5 whole plants for each biological replicate) were either examined immediately or stored at −80°C for future analysis.

For PG isolation, approximately 70 plants were used as starting materials for each biological replicate, following a method adapted from (Espinoza-Corral et al. 2019). The washed chloroplasts were normalized by chlorophyll concentration (∼ 2 mg/ml for control samples, ∼ 1 mg/ml for infected samples), resuspended in an osmotic stress buffer (10 mM Tricine, pH 7.9, 1 mM EDTA, 0.6 M sucrose), and incubated in darkness on ice for 30 minutes. The chloroplasts were stored at −80°C for at least 18 h until the next step. After thawing the chloroplast, the broken chloroplasts were centrifuged at 100000 x *g* for 1 h and 4°C to separate the stroma from the membranes. Chloroplast membranes were resuspended in 48% (w/v) sucrose in HE buffer (50 mM HEPES, pH 7.9, 2 mm EDTA, and 1× cOmplete™ Protease Inhibitor Cocktail (Roche Diagnostics, Mannheim, Germany)) and sonicated with 4 pulses of 10 sec and 20% wattage (Model VCX 130, Sonics & Materials INC, Newtown, CT, USA). Sonicated chloroplast membranes were overlaid with 5% (w/v) sucrose in HE buffer (30 mM HEPES-KOH, pH 8.0, 2 mM EDTA) and centrifuged at 150000 x *g* for 1.5 h and 4°C. PGs (the yellow layer) were collected from the surface of the sucrose gradient.

### Transient expression of proteins via Agrobacterium infiltration

The vector was transformed into *Agrobacterium tumefaciens* GV3101 via electroporation. The *A. tumefaciens* transformants were grown in 5 mL liquid LB with appropriate antibiotics overnight at 28°C, pelleted, and resuspended in induction media (10 mM MES-NaOH, pH 5.6, 10 mM MgCl2, 150 μM acetosyringone) to an OD600 of 0.4–0.6 for transient expression, and incubated in induction media for approximately 3–4 h before infiltration in 5-week-old *N. benthamiana* leaves, which were then incubated at 25°C for 48–72 hours (16 h light/8 h dark) for further analysis.

### Confocal microscopy imaging

All fluorescence detection by confocal microscopy was performed using the Zeiss LSM710 confocal microscope (Carl Zeiss Inc, White Plains, NY, USA) at the Rausser College of Natural Resources Biological Imaging Facility, UC Berkeley.

The 35S:DGAT3-GFP construct was previously described (Xue et al. 2024)). To generate the 35S:FBN1b-mScarlet construct, two BsaI restriction enzyme sites were added to both 5’ and 3’ of the full-length cDNA encoding FBN1b (AT4G22240) without stop codon and the sequence of mScarlet with stop codon using PCR primers listed in **Supplemental Table S1**. Sequences of FBN1b and mScarlet were assembled into the pICSL22011 plasmid under a 35S promoter and 35S terminator via Golden Gate cloning. Subcellular colocalization assays for DGAT3-GFP and FBN1b-mScarlet were performed by transiently co-expressing the proteins in *N. benthamiana* leaves via *Agrobacterium* infiltration. The fluorescence was observed at 48–72 hours post infiltration (hpi). Excitation of GFP and mScarlet were at 488 and 543 nm, respectively. Emission wavelengths for GFP and mScarlet were 493-579 and 582-657 nm, respectively. Excitation and emission wavelengths for chlorophyll were at 633 and 657-721 nm, respectively. The intensity profiling is plotted using Imaris software. The infiltration events were repeated on three different days. The total number of cells observed is 10 and all 10 cells have spots of overlapping GFP and mScarlet signals.

The GFP signals of 35S:DGAT3-GFP in transgenic *Arabidopsis* leaf tissues under untreated, or 3-day darkness incubated (3DDI), or 12dpi conditions, were examined by randomly selecting 10 regions for each treatment. All of these regions showed consistent observations. The GFP signals of 35S:DGAT3-GFP in isolated chloroplasts extracted from transgenic *Arabidopsis* leaf tissues after 3DDI treatment were examined. Approximately 200 chloroplasts were observed, with GFP signals detected in about 15% of them and all GFP signals exhibited a speckled pattern. The GFP signals of 35S:DGAT3-GFP in isolated PGs extracted from transgenic *Arabidopsis* leaf tissues after 3DDI treatment were examined and the total number of PGs observed is around 100. About 50% of PGs show GFP expression and all of those had overlapping fluorescence from Nile Red. Excitation and emission wavelengths of GFP were at 488 and 493-556 nm, respectively.

Excitation and emission wavelengths of Nile Red were at 514 and 541-753 nm, respectively. Golor4_2520209 is annotated in Mycocosm (*Golovinomyces orontii MGH1* v4.0). The signal peptide of Golor4_2520209SC was identified by using SignalP5.0 (https://services.healthtech.dtu.dk/services/SignalP-5.0/) (Almagro Armenteros et al. 2019). The chloroplast transit peptide was determined by LOCALIZER (http://localizer.csiro.au/index.html) (Sperschneider et al. 2017). The transmembrane domain was checked using TMHMM - 2.0 (https://services.healthtech.dtu.dk/services/TMHMM-2.0/) (Krogh et al. 2001). The total RNA of *G. orontii* was extracted from 14 dpi *Arabidopsis* leaves with the RNA Purification Kit (cat no. 74904, QIAGEN Bioinformatics) followed by cDNA synthesis. The cDNA of Golor4_2520209 was amplified using primers listed in **Supplemental Table S1,** and Sanger sequencing showed that a 63 bp region annotated as intron was actually present in the cDNA. Thus, the cDNA, including the 63 bp region, was used for downstream analysis. The cDNA sequence of Golor4_2520209 without the signal peptide was codon-optimized and synthesized by Twist Bioscience (San Francisco, CA, USA). The synthesized Golor4_2520209ΔSP without the stop codon was cloned into the pICSL22010 plasmid under a 35S promoter with a C-terminal GFP and 35S terminator. The transient expression of 35S:Golor4_2520209-GFP in *N. benthamiana* leaf tissues was observed using the same microscope at 48–72 hpi, and the 35S:GFP was used as a negative control. Three independent infiltrations were done. The total number of cells observed is 15, and all 15 cells showed the same localization.

### GFP fluorescence quantification

Two leaf punches were collected from an individual plant at 4–5 weeks old and eight leaf punches in total were used for each genotype per treatment. Each leaf punch was overlaid on top of 300 µL tap water in a well in the 96-well flat bottom with low evaporation lid Microtest tissue culture plate (Becton Dickinson Labware, NJ, USA). Fluorescence measurements were collected with an Infinite M1000 Pro plate-reading spectrophotometer/fluorometer (Tecan, Männedorf, Switzerland). The fluorescence was measured with the following settings: Mode = Fluorescence Top Reading, Multiple Reads per Well (Square (filled)) = 3 x 3, Multiple Reads per Well (Border) = 600 µm, Excitation Wavelength = 488 nm, Emission Wavelength = 510 nm, Excitation Bandwidth = 8 nm, Emission Bandwidth = 8 nm, Gain = 75, Manual Number of Flashes = 50, Flash Frequency = 400 Hz, Integration Time = 20 µs, Lag Time = 0 µs, Settle Time = 10 ms, Z-Position (Calculated From: A1) = 24263 µm. The fluorescence measurement of GFP in transgenic plants was standardized by subtracting the fluorescence measurement of wild plants carrying no transgene. Means with the different letters are significantly different according to one-way ANOVA followed by post-hoc Tukey test (p < 0.05). The whole experiment was repeated twice.

### Y2H assay

The Y2H assay was performed following the steps in the Matchmaker Gold Yeast Two-Hybrid System (Takara Bio USA, Inc., San Jose, CA, USA) manual. The normalized *Arabidopsis* cDNA library was purchased from Takara Bio (cat. no. 630487). The coding region of *Golor4_2520209* without the signal peptide (*Golor4_2520209ΔSP*) was cloned into the *pGBKT7* vector via Golden Gate cloning to generate the construct *BD-Golor4_252020*9 as bait. After mating, colonies were selected on SD/-Leu/-Trp/-Ade/-His DO (QDO) medium after growth at 30°C for 1 week. The prey proteins in these selected colonies were identified by amplifying the DNA sequence using primers listed in **Supplemental Table S1** and Sanger sequencing. To confirm the protein interactions from the screening, the bait, and the prey were co-transformed into yeast cells (strain AH109) and cultured on SD/-Leu/-Trp DO medium. After growth at 30°C for 72 h, independent colonies were selected on SD/-Leu/-Trp/-Ade/-His DO (QDO) medium. The *pGADT7-*empty*/pGBKT7-*empty was used as a negative control. The colonies harboring the target constructs were verified by PCR amplification of both genes.

### Co-IP assay

The coding region of *FBN2 PAP_fibrillin* domain was cloned into the *pICSL22011* vector under a 35S promoter, a C-terminal FLAG-tag and 35S terminator via Golden Gate cloning. To confirm the Golor4_2520209–FBN2 interaction, the OEWSC-GFP and/or FBN2 PAP_fibrillin-Flag, and P19 construct was co-infiltrated into *N.benthamiana* leaves. *N.benthamiana* leaf tissues were harvested at 48-72 hpi and ground on ice in extraction buffer containing 50 mM Tris-HCl (pH 7.4), 1 mM EDTA, 150 mM NaCl, 0.1% (v/v) Triton X-100, 1 mM PMSF, and 1× cOmplete™ Protease Inhibitor Cocktail (Roche Diagnostics, Mannheim, Germany). The extract was centrifuged at 12000 rpm for 2 minutes, and the supernatant was transferred into a fresh tube for the co-IP assay. Immunoprecipitation experiments were performed with µMACS™ GFP Isolation Kit following the manufacturer’s protocol (cat. no. 130-091-125; Miltenyi Biotec, San Diego, CA, USA). In brief, cell lysates were incubated with the anti-GFP coupled magnetic beads for 30 minutes. After running the lysates through the column, the beads were washed four times extensively with 200 µl extraction buffer and the eluted protein was then detected by immunoblotting on the PVDF membrane using an anti-FLAG HRP antibody (1:10,000; cat. no. MA1-91878-HRP, Thermo Fisher Scientific, Waltham, MA, USA) and a mouse anti-GFP (1:5000; cat. no. 11814460001, Roche)) with a secondary goat anti-mouse IgG (whole molecule)-peroxidase-conjugate antibody (1:2000; cat. no. A9044-2ML, Sigma-Aldrich).

### Split-luciferase complementation assay

The coding region of FBN2 PAP_fibrillin domain and Golor4_2520209 without the signal peptide (Golor4_2520209ΔSP) were PCR amplified with primers listed in **Supplemental Table S1**, and digested with KpnI and SalI, and cloned into the nLUC or cLUC vectors. The infiltrated *N. benthamina* leaves expressing split-luciferase complementation constructs at 48–72 hpi were sprayed with 1 mg/ml D-luciferin (cat. no. L-8220, Biosynth Chemistry & Biology) and incubated in darkness for 15 minutes before imaging. The luciferase activities that indicate protein‒protein interactions were detected by a BioRad CCD imaging system (ChemiDoc MP, Model No. Universal Hood III, BioRad). The nLUC/ cLUC plasmid was used as a negative control.

## AUTHOR CONTRIBUTIONS

MCW directed the research. HX, KKN and MCW planned and designed experiments. HX, CK and EC performed experiments, with expertise from MI. HX and MCW wrote the manuscript. All authors contributed to the review of the manuscript.

## ACKNOWLEDGEMENTS

This research was supported by awards to MCW from the American Vineyard Foundation and the Fred C. Gloeckner Foundation. HX also received a NRAEF Viticulture and Enology Scholarship. MI is supported by the US Department of Energy Office of Science through the

Photosynthetic Systems program in the Office of Basic Energy Sciences. KKN is an investigator of the Howard Hughes Medical Institute. We thank Dr. Steven Ruzin and Dr. Denise Schichnes of the RCNR Biological Imaging Facility (UC Berkeley) for assistance with confocal microscopy; this facility is supported in part by the National Institutes of Health S10 program, award number 1S10RR026866-01. We thank Dr. Peter Dörmann (University of Bonn) for providing the *pes1 pes2* double mutant seeds and critical review of the manuscript. We thank Dr. Ángel Mérida (Institute of Plant Biochemistry and Photosynthesis, CSIC-US) for providing the seeds of the *fbn1a-1b* double mutant, *fbn1a-1b-2* triple mutant and *fbn2* complemented lines.

## SUPPORTING INFORMATION

**Supplementary Figure S1.** A zoom-in view of stacked thylakoid membranes separated into the bottom layer of a discontinuous sucrose gradient by centrifugation, supports Figure 2.

**Supplementary Figure S2.** Details of western blots of marked constructs and proteins eluted from anti-GFP coupled magnetic beads detected with the anti-GFP or anti-FLAG antibody, supports Figure 6.

**Supplemental Table S1.** Predicted chloroplast-targeted *G. orontii* effectors

**Supplemental Table S2.** Genotyping, cloning, and SIGS dsRNA template primers used for this work.

**Supplemental Dataset S1**. Yeast-two-hybrid screening results of Golor4_2520209

## DATA AVAILABILITY STATEMENT

All relevant data can be found within the manuscript and its supporting materials.

## REFERENCES

Almagro Armenteros, José Juan, Konstantinos D. Tsirigos, Casper Kaae Sønderby, Thomas Nordahl Petersen, Ole Winther, Søren Brunak, Gunnar von Heijne, and Henrik Nielsen. 2019. “SignalP 5.0 Improves Signal Peptide Predictions Using Deep Neural Networks.” Nature Biotechnology 37 (4): 420–23.

Arzac, Miren I., Beatriz Fernández-Marín, and José I. García-Plazaola. 2022. “More than Just Lipid Balls: Quantitative Analysis of Plastoglobule Attributes and Their Stress-Related Responses.” Planta 255 (3): 62.

Austin, Jotham R., 2nd, Elizabeth Frost, Pierre-Alexandre Vidi, Felix Kessler, and L. Andrew Staehelin. 2006. “Plastoglobules Are Lipoprotein Subcompartments of the Chloroplast That Are Permanently Coupled to Thylakoid Membranes and Contain Biosynthetic Enzymes.” The Plant Cell 18 (7): 1693–1703.

Aymé, Laure, Simon Arragain, Michel Canonge, Sébastien Baud, Nadia Touati, Ornella Bimai, Franjo Jagic, et al. 2018. “Arabidopsis Thaliana DGAT3 Is a [2Fe-2S] Protein Involved in TAG Biosynthesis.” Scientific Reports 8 (1): 1–10.

Bhattacharyya, Dhriti, and Supriya Chakraborty. 2018. “Chloroplast: The Trojan Horse in Plant-Virus Interaction.” Molecular Plant Pathology 19 (2): 504–18.

Bhattacharyya, Dhriti, Prabu Gnanasekaran, Reddy Kishore Kumar, Nirbhay Kumar Kushwaha, Veerendra Kumar Sharma, Mohd Aslam Yusuf, and Supriya Chakraborty. 2015. “A Geminivirus Betasatellite Damages the Structural and Functional Integrity of Chloroplasts Leading to Symptom Formation and Inhibition of Photosynthesis.” Journal of Experimental Botany 66 (19): 5881–95.

Bhuiyan, Nazmul H., Giulia Friso, Elden Rowland, Kristina Majsec, and Klaas J. van Wijk. 2016. “The Plastoglobule-Localized Metallopeptidase PGM48 Is a Positive Regulator of Senescence in Arabidopsis Thaliana.” The Plant Cell 28 (12): 3020–37.

Caplan, Jeffrey L., Amutha Sampath Kumar, Eunsook Park, Meenu S. Padmanabhan, Kyle Hoban, Shannon Modla, Kirk Czymmek, and Savithramma P. Dinesh-Kumar. 2015. “Chloroplast Stromules Function during Innate Immunity.” Developmental Cell 34 (1): 45–57.

Chandran, Divya, Noriko Inada, Greg Hather, Christiane K. Kleindt, and Mary C. Wildermuth. 2010. “Laser Microdissection of Arabidopsis Cells at the Powdery Mildew Infection Site Reveals Site-Specific Processes and Regulators.” Proceedings of the National Academy of Sciences of the United States of America 107 (1): 460–65.

Chandran, Divya, Yu Chuan Tai, Gregory Hather, Julia Dewdney, Carine Denoux, Diane G. Burgess, Frederick M. Ausubel, Terence P. Speed, and Mary C. Wildermuth. 2009. “Temporal Global Expression Data Reveal Known and Novel Salicylate-Impacted Processes and Regulators Mediating Powdery Mildew Growth and Reproduction on Arabidopsis.” Plant Physiology 149 (3): 1435–51.

Channarayappa, Channarayappa, V. Muniyappa, D. Schwegler-Berry, and G. Shivashankar. 1992. “Ultrastructural Changes in Tomato Infected with Tomato Leaf Curl Virus, a Whitefly-Transmitted Geminivirus.” Canadian Journal of Botany. Journal Canadien de Botanique 70 (9): 1747–53.

Chen, Hsu-Ching, Annelyse Klein, Minghui Xiang, Ralph A. Backhaus, and Marcel Kuntz. 1998. “Drought- and Wound-Induced Expression in Leaves of a Gene Encoding a Chromoplast Carotenoid-Associated Protein.” The Plant Journal 14 (3): 317–26.

Clough, S. J., and A. F. Bent. 1998. “Floral Dip: A Simplified Method for Agrobacterium-Mediated Transformation of Arabidopsis Thaliana.” The Plant Journal: For Cell and Molecular Biology 16 (6): 735–43.

Cooper, Bret, Joseph D. Clarke, Paul Budworth, Joel Kreps, Don Hutchison, Sylvia Park, Sonia Guimil, et al. 2003. “A Network of Rice Genes Associated with Stress Response and Seed Development.” Proceedings of the National Academy of Sciences of the United States of America 100 (8): 4945–50.

El-Sappah, Ahmed H., Jia Li, Kuan Yan, Chaoyang Zhu, Qiulan Huang, Yumin Zhu, Yu Chen, Khaled A. El-Tarabily, and Synan F. AbuQamar. 2024. “Fibrillin Gene Family and Its Role in Plant Growth, Development, and Abiotic Stress.” Frontiers in Plant Science 15 (October). 10.3389/fpls.2024.1453974.

Espinoza-Corral, Roberto, Steffen Heinz, Andreas Klingl, Peter Jahns, Martin Lehmann, Jörg Meurer, Jörg Nickelsen, Jürgen Soll, and Serena Schwenkert. 2019. “Plastoglobular Protein 18 Is Involved in Chloroplast Function and Thylakoid Formation.” Journal of Experimental Botany 70 (15): 3981–93.

Espinoza-Corral, Roberto, Serena Schwenkert, and Peter K. Lundquist. 2021. “Molecular Changes of Arabidopsis Thaliana Plastoglobules Facilitate Thylakoid Membrane Remodeling under High Light Stress.” The Plant Journal: For Cell and Molecular Biology 106 (6): 1571–87.

Froehlich, John E., Curtis G. Wilkerson, W. Keith Ray, Rosemary S. McAndrew, Katherine W. Osteryoung, Douglas A. Gage, and Brett S. Phinney. 2003. “Proteomic Study of the Arabidopsis Thaliana Chloroplastic Envelope Membrane Utilizing Alternatives to Traditional Two-Dimensional Electrophoresis.” Journal of Proteome Research 2 (4): 413–25.

Gámez-Arjona, Francisco Manuel, Juan Carlos de la Concepción, Sandy Raynaud, and Ángel Mérida. 2014. “Arabidopsis Thaliana Plastoglobule-Associated Fibrillin 1a Interacts with Fibrillin 1b in Vivo.” FEBS Letters 588 (17): 2800–2804.

Gámez-Arjona, Francisco M., Sandy Raynaud, Paula Ragel, and Angel Mérida. 2014. “Starch Synthase 4 Is Located in the Thylakoid Membrane and Interacts with Plastoglobule-Associated Proteins in Arabidopsis.” The Plant Journal: For Cell and Molecular Biology 80 (2): 305–16.

Gaude, Nicole, Claire Bréhélin, Gilbert Tischendorf, Felix Kessler, and Peter Dörmann. 2007. “Nitrogen Deficiency in Arabidopsis Affects Galactolipid Composition and Gene Expression and Results in Accumulation of Fatty Acid Phytyl Esters.” The Plant Journal: For Cell and Molecular Biology 49 (4): 729–39.

Jelenska, Joanna, Nan Yao, Boris A. Vinatzer, Christine M. Wright, Jeffrey L. Brodsky, and Jean T. Greenberg. 2007. “A J Domain Virulence Effector of Pseudomonas Syringae Remodels Host Chloroplasts and Suppresses Defenses.” Current Biology: CB 17 (6): 499–508.

Jiang, Er-Ying, Yong Fan, Nghi-Van Phung, Wan-Yue Xia, Guang-Rong Hu, and Fu-Li Li. 2023. “Overexpression of Plastid Lipid-Associated Protein in Marine Diatom Enhances the Xanthophyll Synthesis and Storage.” Frontiers in Microbiology 14 (April):1143017.

Jiang, Yina, Wanxiao Wang, Qiujin Xie, Na Liu, Lixia Liu, Dapeng Wang, Xiaowei Zhang, et al. 2017. “Plants Transfer Lipids to Sustain Colonization by Mutualistic Mycorrhizal and Parasitic Fungi.” Science 356 (6343): 1172–75.

Kanwischer, Marion, Svetlana Porfirova, Eveline Bergmüller, and Peter Dörmann. 2005. “Alterations in Tocopherol Cyclase Activity in Transgenic and Mutant Plants of Arabidopsis Affect Tocopherol Content, Tocopherol Composition, and Oxidative Stress.” Plant Physiology 137 (2): 713–23.

Kim, Inyoung, Eun-Ha Kim, Yu-Ri Choi, and Hyun Uk Kim. 2022. “Fibrillin2 in Chloroplast Plastoglobules Participates in Photoprotection and Jasmonate-Induced Senescence.” Plant Physiology 189 (3): 1363–79.

Krogh, A., B. Larsson, G. von Heijne, and E. L. Sonnhammer. 2001. “Predicting Transmembrane Protein Topology with a Hidden Markov Model: Application to Complete Genomes.” Journal of Molecular Biology 305 (3): 567–80.

Laizet, Yechan, Dominique Pontier, Régis Mache, and Marcel Kuntz. 2004. “Subfamily Organization and Phylogenetic Origin of Genes Encoding Plastid Lipid-Associated Proteins of the Fibrillin Type.” Journal of Genome Science and Technology 3 (1): 19–28.

Lee, Mi Yeon, Johan Jaenisch, Amanda G. McRae, Chihiro Hirai-Adachi, Katherine Louie, Leslie P. Silva, Benjamin P. Bowen, Trent R. Northen, and Mary C. Wildermuth. 2024. “Altered Host Pyruvate Metabolism Fuels and Regulates Fungal Asexual Reproduction.” bioRxiv, January, 2024.08.22.609210.

Li, Jiajia, Xukai Li, Ahmed Adel Khatab, and Guosheng Xie. 2020. “Phylogeny, Structural Diversity and Genome-Wide Expression Analysis of Fibrillin Family Genes in Rice.” Phytochemistry 175 (112377): 112377.

Lippold, Felix, Katharina vom Dorp, Marion Abraham, Georg Hölzl, Vera Wewer, Jenny Lindberg Yilmaz, Ida Lager, et al. 2012. “Fatty Acid Phytyl Ester Synthesis in Chloroplasts of Arabidopsis.” The Plant Cell 24 (5): 2001–14.

Littlejohn, George R., Susan Breen, Nicholas Smirnoff, and Murray Grant. 2021. “Chloroplast Immunity Illuminated.” The New Phytologist 229 (6): 3088–3107.

Liu, Jie, Pan Gong, Ruobin Lu, Rosa Lozano-Durán, Xueping Zhou, and Fangfang Li. 2024. “Chloroplast Immunity: A Cornerstone of Plant Defense.” Molecular Plant 17 (5): 686–88.

Li, Xiao, Yuhan Liu, Qiguang He, Sipeng Li, Wenbo Liu, Chunhua Lin, and Weiguo Miao. 2020. “A Candidate Secreted Effector Protein of Rubber Tree Powdery Mildew Fungus Contributes to Infection by Regulating Plant ABA Biosynthesis.” Frontiers in Microbiology 11 (November):591387.

Lohscheider, Jens N., Giulia Friso, and Klaas J. van Wijk. 2016. “Phosphorylation of Plastoglobular Proteins in Arabidopsis Thaliana.” Journal of Experimental Botany 67 (13): 3975–84.

Lundquist, Peter K., Anton Poliakov, Nazmul H. Bhuiyan, Boris Zybailov, Qi Sun, and Klaas J. van Wijk. 2012. “The Functional Network of the Arabidopsis Plastoglobule Proteome Based on Quantitative Proteomics and Genome-Wide Coexpression Analysis.” Plant Physiology 158 (3): 1172–92.

Lundquist, Peter K., Anton Poliakov, Lisa Giacomelli, Giulia Friso, Mason Appel, Ryan P. McQuinn, Stuart B. Krasnoff, et al. 2013. “Loss of Plastoglobule Kinases ABC1K1 and ABC1K3 Causes Conditional Degreening, Modified Prenyl-Lipids, and Recruitment of the Jasmonic Acid Pathway.” The Plant Cell 25 (5): 1818–39.

Lu, Yan, and Jian Yao. 2018. “Chloroplasts at the Crossroad of Photosynthesis, Pathogen Infection and Plant Defense.” International Journal of Molecular Sciences 19 (12). 10.3390/ijms19123900.

Mandal, Md Siddikun Nabi, Ying Fu, Sheng Zhang, and Wanquan Ji. 2014. “Proteomic Analysis of the Defense Response of Wheat to the Powdery Mildew Fungus, Blumeria Graminis F. Sp. Tritici.” The Protein Journal 33 (6): 513–24.

McRae, Amanda G., Jyoti Taneja, Kathleen Yee, Xinyi Shi, Sajeet Haridas, Kurt LaButti, Vasanth Singan, Igor V. Grigoriev, and Mary C. Wildermuth. 2023. “Spray-Induced Gene Silencing to Identify Powdery Mildew Gene Targets and Processes for Powdery Mildew Control.” *Molecular Plant Pathology*, June. 10.1111/mpp.13361.

Micali, Cristina, Katharina Göllner, Matt Humphry, Chiara Consonni, and Ralph Panstruga. 2008. “The Powdery Mildew Disease of Arabidopsis: A Paradigm for the Interaction between Plants and Biotrophic Fungi.” The Arabidopsis Book / American Society of Plant Biologists 6 (October):e0115.

Mu, Bo, Zhaolin Teng, Ruixin Tang, Mengjiao Lu, Jinfu Chen, Xiangnan Xu, and Ying-Qiang Wen. 2023. “An Effector of Erysiphe Necator Translocates to Chloroplasts and Plasma Membrane to Suppress Host Immunity in Grapevine.” Horticulture Research 10 (9): uhad163.

Norkunas, Karlah, Robert Harding, James Dale, and Benjamin Dugdale. 2018. “Improving Agroinfiltration-Based Transient Gene Expression in Nicotiana Benthamiana.” Plant Methods 14 (1): 71.

Otulak, Katarzyna, Marcin Chouda, Józef Bujarski, and Grażyna Garbaczewska. 2015. “The Evidence of Tobacco Rattle Virus Impact on Host Plant Organelles Ultrastructure.” Micron (Oxford, England: 1993) 70 (March):7–20.

Prokopová, Jitka, Martina Spundová, Michaela Sedlárová, Alexandra Husicková, Radko Novotný, Karel Dolezal, Jan Naus, and Ales Lebeda. 2010. “Photosynthetic Responses of Lettuce to Downy Mildew Infection and Cytokinin Treatment.” Plant Physiology and Biochemistry: PPB / Societe Francaise de Physiologie Vegetale 48 (8): 716–23.

Reuber, T. L., J. M. Plotnikova, J. Dewdney, E. E. Rogers, W. Wood, and F. M. Ausubel. 1998. “Correlation of Defense Gene Induction Defects with Powdery Mildew Susceptibility in Arabidopsis Enhanced Disease Susceptibility Mutants.” The Plant Journal: For Cell and Molecular Biology 16 (4): 473–85.

Rey, P., B. Gillet, S. Römer, F. Eymery, J. Massimino, G. Peltier, and M. Kuntz. 2000. “Over-Expression of a Pepper Plastid Lipid-Associated Protein in Tobacco Leads to Changes in Plastid Ultrastructure and Plant Development upon Stress.” The Plant Journal: For Cell and Molecular Biology 21 (5): 483–94.

Rodríguez-Herva, José J., Pablo González-Melendi, Raquel Cuartas-Lanza, María Antúnez-Lamas, Isabel Río-Alvarez, Ziduo Li, Gema López-Torrejón, et al. 2012. “A Bacterial Cysteine Protease Effector Protein Interferes with Photosynthesis to Suppress Plant Innate Immune Responses: P. Syringae HopN1 Targets Photosynthesis.” Cellular Microbiology 14 (5): 669–81.

Rottet, Sarah, Julie Devillers, Gaétan Glauser, Véronique Douet, Céline Besagni, and Felix Kessler. 2016. “Identification of Plastoglobules as a Site of Carotenoid Cleavage.” Frontiers in Plant Science 7 (December):1855.

Shanmugabalaji, Venkatasalam, Bernhard Grimm, and Felix Kessler. 2020. “Characterization of a Plastoglobule-Localized SOUL4 Heme-Binding Protein in Arabidopsis Thaliana.” Frontiers in Plant Science 11 (January):2.

Singh, Dharmendra K., Siela N. Maximova, Philip J. Jensen, Brian L. Lehman, Henry K. Ngugi, and Timothy W. McNellis. 2010. “FIBRILLIN4 Is Required for Plastoglobule Development and Stress Resistance in Apple and Arabidopsis.” Plant Physiology 154 (3): 1281–93.

Singh, Dharmendra K., and Timothy W. McNellis. 2011. “Fibrillin Protein Function: The Tip of the Iceberg?” Trends in Plant Science 16 (8): 432–41.

Sperschneider, Jana, Ann-Maree Catanzariti, Kathleen DeBoer, Benjamin Petre, Donald M. Gardiner, Karam B. Singh, Peter N. Dodds, and Jennifer M. Taylor. 2017. “LOCALIZER: Subcellular Localization Prediction of Both Plant and Effector Proteins in the Plant Cell.” Scientific Reports 7 (March):44598.

Tominaga, Jun, Yasutoshi Nakahara, Daisuke Horikawa, Ayumi Tanaka, Maki Kondo, Yasuhiro Kamei, Tsuneaki Takami, et al. 2018. “Overexpression of the Protein Disulfide Isomerase AtCYO1 in Chloroplasts Slows Dark-Induced Senescence in Arabidopsis.” BMC Plant Biology 18 (1): 80.

Torres-Romero, Diego, Ángeles Gómez-Zambrano, Antonio Jesús Serrato, Mariam Sahrawy, and Ángel Mérida. 2022. “Arabidopsis Fibrillin 1-2 Subfamily Members Exert Their Functions via Specific Protein-Protein Interactions.” Journal of Experimental Botany 73 (3): 903–14.

Torres Zabala, Marta de, George Littlejohn, Siddharth Jayaraman, David Studholme, Trevor Bailey, Tracy Lawson, Michael Tillich, et al. 2015. “Chloroplasts Play a Central Role in Plant Defence and Are Targeted by Pathogen Effectors.” Nature Plants 1 (6): 15074.

Wang, Haixia, Ely Oliveira-Garcia, Petra C. Boevink, Nicholas J. Talbot, Paul R. J. Birch, and Barbara Valent. 2023. “Filamentous Pathogen Effectors Enter Plant Cells via Endocytosis.” Trends in Plant Science 28 (11): 1214–17.

Weßling, Ralf, Petra Epple, Stefan Altmann, Yijian He, Li Yang, Stefan R. Henz, Nathan McDonald, et al. 2014. “Convergent Targeting of a Common Host Protein-Network by Pathogen Effectors from Three Kingdoms of Life.” Cell Host & Microbe 16 (3): 364–75.

Wijk, Klaas J. van, and Felix Kessler. 2017. “Plastoglobuli: Plastid Microcompartments with Integrated Functions in Metabolism, Plastid Developmental Transitions, and Environmental Adaptation.” Annual Review of Plant Biology 68 (April):253–89.

Xu, Qiang, Chunlei Tang, Xiaodong Wang, Shutian Sun, Jinren Zhao, Zhensheng Kang, and Xiaojie Wang. 2019. “An Effector Protein of the Wheat Stripe Rust Fungus Targets Chloroplasts and Suppresses Chloroplast Function.” Nature Communications 10 (1): 5571.

Xue, Hang, Johan Jaenisch, Joelle Sasse, Erin Riley McGarrigle, Emma H. Choi, Katherine B. Louie, Katharina Gutbrod, Peter Dörmann, Trent Northen, and Mary C. Wildermuth. 2024. “Powdery Mildew Infection Induces a Novel Route to Storage Lipid Formation at the Expense of Host Thylakoid Lipids to Promote Fungal Spore Production.” The Plant Cell, in press.

Yang, Feng, Kunqin Xiao, Hongyu Pan, and Jinliang Liu. 2021. “Chloroplast: The Emerging Battlefield in Plant–Microbe Interactions.” Frontiers in Plant Science 12. 10.3389/fpls.2021.637853.

Yang, Yi, Ronan Sulpice, Axel Himmelbach, Michael Meinhard, Alexander Christmann, and Erwin Grill. 2006. “Fibrillin Expression Is Regulated by Abscisic Acid Response Regulators and Is Involved in Abscisic Acid-Mediated Photoprotection.” Proceedings of the National Academy of Sciences of the United States of America 103 (15): 6061–66.

Youssef, Abir, Yec ’han Laizet, Maryse A. Block, Eric Maréchal, Jean-Pierre Alcaraz, Tony R. Larson, Dominique Pontier, Joël Gaffé, and Marcel Kuntz. 2010. “Plant Lipid-Associated Fibrillin Proteins Condition Jasmonate Production under Photosynthetic Stress.” The Plant Journal: For Cell and Molecular Biology 61 (3): 436–45.

Ytterberg, A. Jimmy, Jean-Benoit Peltier, and Klaas J. van Wijk. 2006. “Protein Profiling of Plastoglobules in Chloroplasts and Chromoplasts. A Surprising Site for Differential Accumulation of Metabolic Enzymes.” Plant Physiology 140 (3): 984–97.

Zechmann, Bernd. 2019. “Ultrastructure of Plastids Serves as Reliable Abiotic and Biotic Stress Marker.” PloS One 14 (4): e0214811.

Zhang, Quancheng, Menghan Zhou, and Jungang Wang. 2022. “Increasing the Activities of Protective Enzymes Is an Important Strategy to Improve Resistance in Cucumber to Powdery Mildew Disease and Melon Aphid under Different Infection/infestation Patterns.” Frontiers in Plant Science 13 (August):950538.

