## Supplemental Information for "Powdery mildew exploits host plastoglobuli functions via DGAT3 and FBN2 for proliferation"

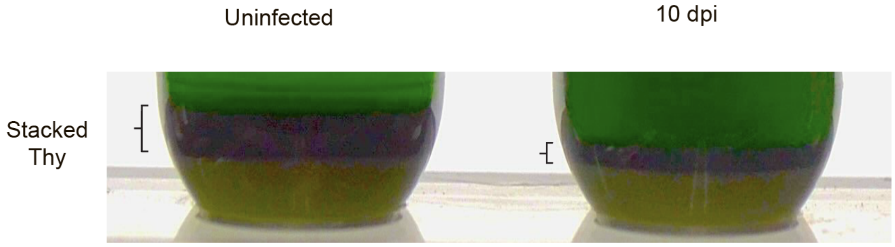


**Supplementary Figure S1. A zoom-in view of stacked thylakoid separated into the bottom layer of a discontinuous sucrose gradient by centrifugation, supports Figure 2.**

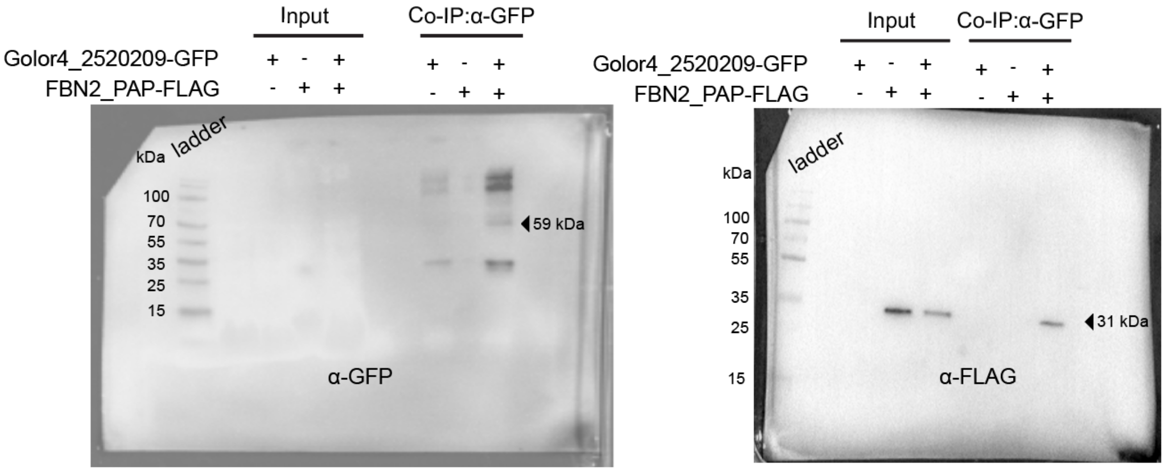


**Supplementary Figure S2. Details of western blots of marked constructs and proteins eluted from anti-GFP coupled magnetic beads detected with the anti-GFP or anti-FLAG antibody, supports Figure 6.**

**Supplemental Table S1. Predicted chloroplast-targeted *G.orontii* effectors**

| Golor4_ProteinID (and homologs[b]) | SignalP 5.0 | TMHM M2.0 | Localizer | *In planta* localization |
|---|---|---|---|---|
| Golor4_4152115 \| Golor4_4149086 | Yes | No | cTP | cytoplasm and nucleus |
| Golor4_5402089 \| Golor4_4145494 | Yes | No | cTP | cytoplasm |
| Golor4_5249087 | Yes | No | cTP | low expression, inconclusive |
| Golor4_4735914 | Yes | No | -[a] | low expression, inconclusive |
| Golor4_4147523 | Yes | No | cTP | low expression, inconclusive |
| Golor4_2914799 | Yes | No | cTP | cytoplasm and nucleus |
| Golor4_2520209 \| Golor4_1673217 | Yes | No | cTP | chloroplast |
| Golor4_4621148 | Yes | No | cTP | low expression, inconclusive |
| Golor4_5337153 | Yes | No | cTP | cloning unsuccessful |
| Golor4_2908844 \| Golor4_2925102 | Yes | No | cTP | cytoplasm and nucleus |
| Golor4_4149526 | Yes | No | cTP | cloning unsuccessful |
| Golor4_5340314 | Yes | No | cTP | cloning unsuccessful |
| Golor4_2911133 \| Golor4_5328180 | Yes | No | cTP | low expression, inconclusive |
| Golor4_4465299 | Yes | No | cTP | cytoplasm and nucleus |
| Golor4_4746636 \| Golor4_5031231 | Yes | No | cTP | low expression, inconclusive |
| Golor4_5396989 | Yes | No | cTP | cloning unsuccessful |
| Golor4_4153343 \| Golor4_4145820 | Yes | No | cTP | cytoplasm |

a: Although this protein does not have predicted cTP, Weßling et al. identified its ortholog interacts with two chloroplast associated proteins via Y2H (Weßling et al. 2014).

b: Most Golor4 genes have two copies due to recent duplication, with >95% sequence identity.

**Supplemental Table S2. Genotyping, cloning, and SIGS dsRNA template primers used for this work.**

| **Genotyping primers** | | **Purpose** |
|---|---|---|
| G1 | CTATCCAGGCCCATTTAGTCC | *fbn1a-1, SALK_024528C* |
| G2 | TCACCGGGAAATTAAACTTCC | *fbn1a-1, SALK_024528C* |
| G5 | TGTAATCAGCAATCCATTCCG | *fbn2-1, SALK_124590* |
| G6 | TTCAGTTCCGTACACCGAATC | *fbn2-1, SALK_124590* |
| G7 | GGAGGCAAGAAATCAGAAACC | *pes1-1, SALK_034549* |
| G8 | TTGATAGGTCACCGCTACAGC | *pes1-1, SALK_034549* |
| G9 | AGAGGTTGCACCATGAAGTTG | *pes2-1, SALK_071769* |
| G10 | ACTTTCTTGGCTCTTTCTCGG | *pes2-1, SALK_071769* |
| G11 | CAACACAGATGCTTGATGGG | *dgat3-2, SALK_112303* |
| G12 | AGAAGGCTAAAGCCAAAGCC | *dgat3-2, SALK_112303* |
| **SIGS dsRNA template primers** | | |
| SIGS1 | CTTCTGTACGGTGCTAGTTATGGC | Golor4_2520209 SIGS dsRNA F |
| SIGS2 | CGACTGGAGGTAGGGTTGATCC | Golor4_2520209 SIGS dsRNA R |
| SIGS3 | TAATACGACTCACTATAGGGGGCTTCTGTACGGTGCTAGTTATGGC | Golor4_2520209 SIGS dsRNA F + T7 |
| SIGS4 | TAATACGACTCACTATAGGGGGCGACTGGAGGTAGGGTTGATCC | Golor4_2520209 SIGS dsRNA R + T7 |
| **Cloning primers** | | |
| C1 | GCTGCTCTTCGTCTGGTCTCGAATGGCGACGGTACAATTGTCCAC | FBN1b F pICSL22011 |
| C2 | GGCTCTTCATCTGGTCTCGACCTCCAGGATTCAAGAGAGGGCTTCCTTC | FBN1b R pICSL22011 |
| C3 | GCTGCTCTTCGTCTGGTCTCGAGGTATGGTGAGCAAGGGCGAG | mScarlet F pICSL22011 |
| C4 | GGCTCTTCATCTGGTCTCGCGAACCCTTGTACAGCTCGTCCATGCC | mScarlet R pICSL22011 |

| | | |
|---|---|---|
| C5 | GCTGCTCTTCGTCTGGTCTCGAATGGCATCGGCTTCTCCAAA | Golor4_2520209 F pICSL22010 |
| C6 | GGCTCTTCATCTGGTCTCGCGAACCGGCGTTGGTGACCTTTAGTATCC | Golor4_2520209 R pICSL22010 |
| C7 | GCTGCTCTTCGTCTGGTCTCGAATGGCATCGGCTTCTCCAAA | Golor4_2520209 F pGADT7 or pGBKT7 |
| C8 | GGCTCTTCATCTGGTCTCGAAGCCCTTTTAGGCGTTGGTGACCTTTAGTATCC | Golor4_2520209 R pGADT7 or pGBKT7 |
| C9 | GCTGCTCTTCGTCTGGTCTCGAATGGCTACGCTCTTCACCG | FBN2_Full F pGADT7 or pGBKT7 |
| C10 | GGCTCTTCATCTGGTCTCGAAGCCCTCAGAGCTCAAGCAGAGAGCTTC | FBN2_Full R pGADT7 or pGBKT7 |
| C11 | GCTGCTCTTCGTCTGGTCTCGAATGGAGCTGAAAAGGTGTTTG | FBN2_PAP F pGADT7 or pGBKT7 |
| C12 | GGCTCTTCATCTGGTCTCGAAGCCCTCAGAGCTCAAGCAGAGAGCTTC | FBN2_PAP R pGADT7 or pGBKT7 |
| C13 | GCTGCTCTTCGTCTGGTCTCGAATGAGGCTAAAGAGATCACTAG | FBN1a_PAP F pGADT7 or pGBKT7 |
| C14 | GGCTCTTCATCTGGTCTCGAAGCCCTTAAGGGTTTAAGAGAG | FBN1a_PAP R pGADT7 or pGBKT7 |
| C15 | GCTGCTCTTCGTCTGGTCTCGAATGCGGCTAAAGAGATCGCTC | FBN1b_PAP F pGADT7 or pGBKT7 |
| C16 | GGCTCTTCATCTGGTCTCGAAGCCCTCAAGGATTCAAGAGAGG | FBN1b_PAP R pGADT7 or pGBKT7 |
| C17 | GCTGCTCTTCGTCTGGTCTCGAATGGGAAGTGCGGTGTCTGACC | FBN2 F pICSL22011 |
| C18 | GGCTCTTCATCTGGTCTCGCGAACCGAGCTCAAGCAGAGAGCTTCC | FBN2 R pICSL22011 |
| C19 | GGCTCGGTACCGGAGGTATGGAGCTGAAAAGGTGTTTGG | FBN2 F nLUC or cLUC |
| C20 | CTGCAGGTCGACCTCGAGACCGAGCTCAAGCAGAGAGC | FBN2 R nLUC |
| C21 | CTGCAGGTCGACCTCGAGCTAACCGAGCTCAAGCAGAGAGC | FBN2 R cLUC |
| C22 | GGCTCGGTACCGGAGGTATGGCCAGCGCGAGTCCTAAAG | Golor4_2520209 F nLUC or cLUC |
| C23 | CTGCAGGTCGACCTCGAGCGCATTAGTTACTTTAAGAATTC | Golor4_2520209 R nLUC |
| C24 | CTGCAGGTCGACCTCGAGCTACGCATTAGTTACTTTAAGAATTC | Golor4_2520209 R cLUC |
